# *LocalZProjector* and *DeProj*: A toolbox for local 2D projection and accurate morphometrics of large 3D microscopy images

**DOI:** 10.1101/2021.01.15.426809

**Authors:** Sébastien Herbert, Léo Valon, Laure Mancini, Nicolas Dray, Paolo Caldarelli, Jérôme Gros, Elric Esposito, Spencer L. Shorte, Laure Bally-Cuif, Romain Levayer, Nathalie Aulner, Jean-Yves Tinevez

## Abstract

**Background:** Quantitative imaging of epithelial tissues prompts for bioimage analysis tools that are widely applicable and accurate. In the case of imaging 3D tissues, a common pre-processing step consists in projecting the acquired 3D volume on a 2D plane mapping the tissue surface. Indeed, while segmenting the tissue cells is amenable on 2D projections, it is still very difficult and cumbersome in 3D. However, for many specimen and models used in Developmental and Cell Biology, the complex content of the image volume surrounding the epithelium in a tissue often reduces the visibility of the biological object in the projection, compromising its subsequent analysis. In addition, the projection may distort the geometry of the tissue and can lead to strong artifacts in the morphology measurement.

**Results:** Here we introduce a user-friendly toolbox built to robustly project epithelia on their 2D surface from 3D volumes, and to produce accurate morphology measurement corrected for the projection distortion, even for very curved tissues. Our toolbox is built upon two components. *LocalZProjector* is a user-friendly and configurable Fiji plugin that generates 2D projections and height-maps from potentially large 3D stacks (larger than 40 GB per time-point) by only incorporating signal of the planes with local highest variance/mean intensity, despite a possibly complex image content. *DeProj* is a MATLAB tool that generates correct morphology measurements by combining the height-map output (such as the one offered by *LocalZProjector*) and the results of a cell segmentation on the 2D projection, hence effectively deprojecting the 2D segmentation in 3D. In this paper we demonstrate their effectiveness over a wide range of different biological samples. We then compare its performance and accuracy against similar existing tools.

**Conclusions:** We find that *LocalZProjector* performs well even in situations where the volume to project also contains unwanted signal in other layers. We show that it can process large images without a pre-processing step. We study the impact of geometrical distortions on morphological measurements induced by the projection. We measured very large distortions which are then corrected by *DeProj*, providing accurate outputs.

## Background

Tissue morphogenesis and homeostasis emerge from the integration of the properties and behaviours of a large number of cells (from hundreds to millions). This includes the regulation and shaping of epithelia, essential tissues barriers composed of one or several layer of cells tightly bound to one another. The emergence of complex epithelial shape relies quite often on the integration of local mechanical and biochemical cues regulated at the cellular levels (1). For instance, during *Drosophila* pupal development, local cell-to-cell interactions through adhesion forces ensure tissue integrity, control cell shape and cell division orientation and affect tissue-wide cell death probability (2, 3). In the avian embryo during gastrulation, mechanical properties of cells affect the emergent tissue fluidity which is essential to form the first tissue fold (4). In vertebrate adult brains, neural stem cells (NSCs) are organized in a pool forming an epithelium. The architecture and size of the brain relies on the rate of division and differentiation of stem cells which is in part regulated by local cellcell interactions (5). All these examples illustrates an essential challenge of tissue and developmental biology: namely bridging the gap between the cellular scales and emerging properties at the tissue level. The capacity to image large tissues, while reaching a single cell resolution amenable for the quantification of cell scale parameters is key to elucidate this question.

Modern microscopy technologies can fulfill this imaging challenge and microscopes that can acquire fluorescence images of the embryo and generate volumetric, multi-channel, time-lapse datasets of a live sample are now used routinely. They offer single-cell resolution on a field-of-view large enough to encompass a significant part of the tissue/embryo studied. However, the large size of the data generated and the limited image signal-to-noise ratio (SNR) imposed by preservation of the tissue health during live-imaging call for optimized image analysis tools.

Epithelia are a continuous layer of cells forming a smooth surface which may not be flat. When cells are labeled with a junction marker (*e.g*. E-cadherin, ZO1), the tissue resembles a manifold wrapped on a 3D surface. A simple approach to visualization and analysis is to perform a projection of the tissue 3D surface on a 2D plane. This dimensionality-reduction approach proves to be particularly convenient: First the resulting data size is considerably diminished. Second the visualization of the tissue layer content is immediate in 2D. Third, it may enhance signal-to-noise ratio. Finally, most of the segmentation algorithms that can extract cell shapes have a better robustness and accuracy in 2D than in 3D so far, and manual corrections can still be reasonably performed in 2D.

The biological significance of the information extracted from these manifolds prompted for the development of several tools that can perform 2D projection. MorphoGraph-X (6) and ImSAnE (7) belong to a first class of tools, where a reference surface is built as a mesh mapping the sample boundary. The fluorescence intensity is then collected at or a few microns away from the boundary into the sample, in a direction perpendicular to the surface. This approach is particularly adequate for images of samples with complex, possibly closed boundaries. A second class of tools perform a projection of the 3D volume along the Z-axis of the 3D image. The resulting projection created by these tools is a 2D plane that has the same width and height as the source 3D volume. They cannot harness samples where the tissue layer folds, since there must be at most one surface Z value per (X, Y) position. They are however particularly convenient with classical confocal microscopy. Indeed, the point-spread-function (PSF) of a microscope is the most elongated along the Z axis. This causes some larger blurring of structures in this direction compared to along the X and Y axes. By projecting along the Z axis, the projections given by these techniques are devoid of this large blurring. The most simple projection technique in this class consists in taking the largest pixel value along a Z column for each (X, Y) position. As noted in (8), this maximum-intensity projection (*MIP*) technique is the most commonly used by biologists. An important drawback is that this projection incorporates noise from throughout the sample, in particular inside cells, and may compromise segmentation based on the membrane signal. To address this limitation, several projection tools have been developed that aim at including in the projection only the signal coming from the tissue layer. Among them are *StackFocuser* (9), *PreMosa* (10), *Extended Depth of Field* (*EDF*) (11), *SurfCut* (12), *MinCostZSurface* (13–15), the Smooth Manifold Extraction (*SME*) tool (8) and a new implementation of the latter: *FastSME* (16) (see Additional File 1: Supplementary Note 2 for descriptions). In (17), authors also proposed an approach based on Deep-Learning for the projection along the Z-axis, but it requires for its training and along with the source images a possibly large set of images pre-projected by an other method or manually. While these approaches have been proven to work well for tissue imaging, they left open some challenges. First, bright or noisy structures outside of the cell layer might compromise the extraction of a meaningful reference surface, in turn strongly affecting the quality of the projection. Second, most of those approaches will be very time consuming for large volume and may even not be working when the volume to project has a size larger than the available computer memory. Third, the tissue surface and the XY plane may have in some regions a large angle. Beyond 30°, the morphological features of cells measured on the 2D projection will be significantly altered, as noted in (12).

Our toolbox fits into this second class of tools, and builds upon them to address several challenges they left open. *LocalZProjector* is its first component, and is a user-friendly and widely configurable projection tool that can be tuned to detect the right reference surface. Its manoeuvrability makes it amenable to a wide range of samples and image qualities, as well as to images of large size beyond the size of the available computer memory. *DeProj*, its second component, can correct for the distortions induced by the projection of tissue even if they are very curved, hence deprojecting the 2D segmentation results in 3D. We illustrate below these capacities using three different samples: the *Drosophila* pupal notum, the quail embryo and the adult zebrafish telencephalon. We also compare the toolbox features, performance and accuracy to existing tools.

## Implementation

The software toolbox we present is made of two components: *LocalZProjector* and *DeProj* (Figure 1). It is built for accurate morphology measurements on epithelial tissues.

**Fig. 1.**
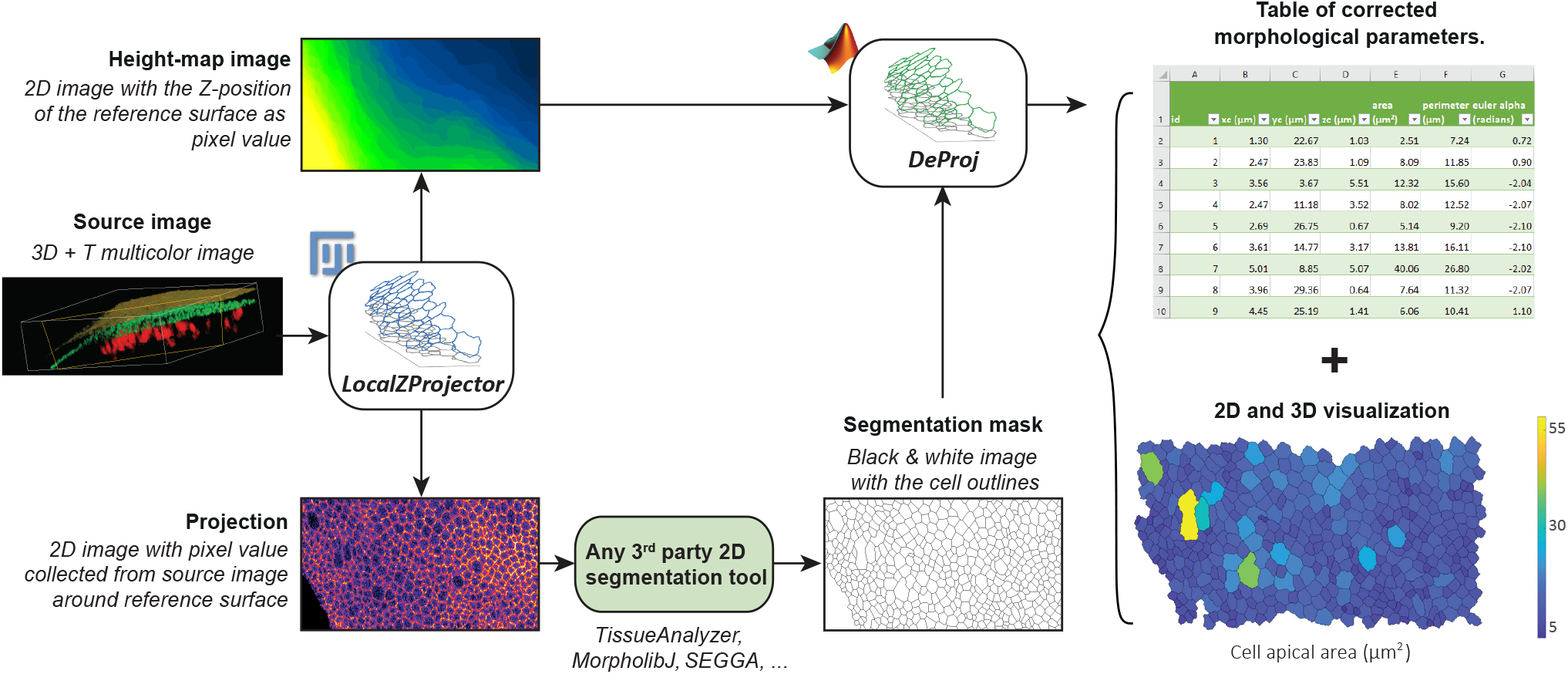
Presentation of *LocalZProjector* and *DeProj*. The toolbox is made of two tools, *LocalZProjector* a Fiji tool that generates 2D projections from 3D, multi-channel time-lapse images, and *DeProj*, a MATLAB function that uses the height-map output of *LocalZProjector* and the segmentation results on the projection output to measure accurately the morphology metrics of the cells in the projected tissue.

*LocalZProjector* performs the projection of a curved surface from a 3D stack on a single 2D plane by only including the signal of the planes with local highest variance/mean intensity, corresponding to the signal of interest (Figure 2a). It is an ImageJ2 (18) plugin distributed within Fiji (19) that focuses on usability and is designed to be adaptable to many different cases and image quality (Figure 2b). It can work with 3D time-lapses, multiple color channels, takes advantage of computers with multiple cores, can be used in scripts and can process images too large to fit in memory using the virtual stack feature of ImageJ. The local Z projection processes as follow: First it extracts a reference surface that maps the epithelial layer (Figure 2c, Additional File 1: Fig S1). The reference surface is represented by the *heightmap*, which consists in a single 2D image per time-point of the source image, that specifies for every (X, Y) position the Z position of the layer of interest. Second, the height-map is used to extract projections of the different channels from the 3D image, according to their relative, sometimes different, offset with this reference surface.

**Fig. 2.**
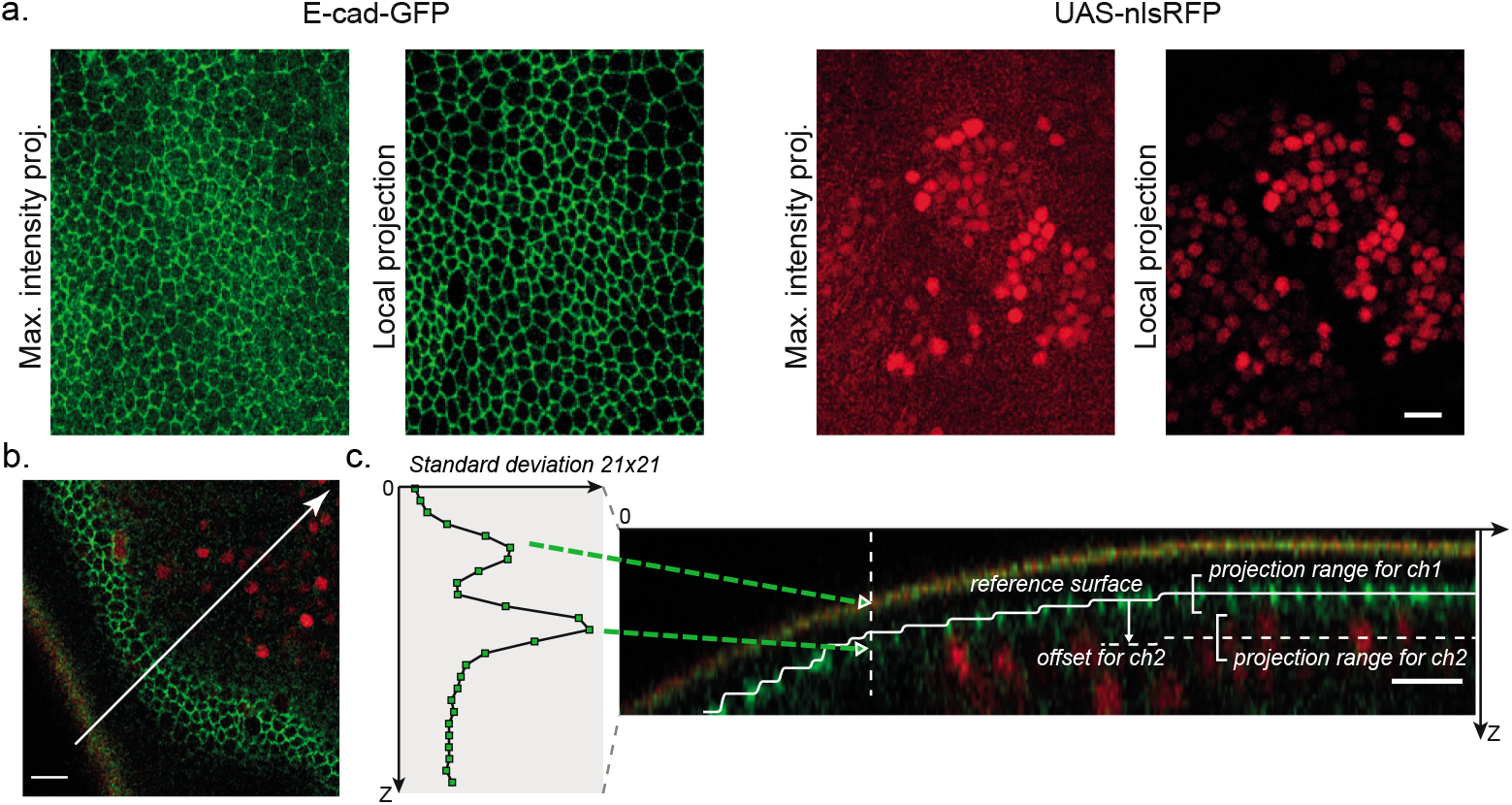
The *LocalZProjector* component. **a.** Comparing the output of the maximum intensity projection (MIP) output with the *LocalZProjector* output on the *Drosophila* pupal notum expressing E-cadherin-GFP. The MIP incorporates signal from other layers (cuticle, fat body) in the projection that compromises its subsequent analysis. The use of a local projection that only includes the signal around a reference surface yields better quality projections. **b.** Example of a single Z-plane from the same dataset showing E-cad::GFP (green) and UAS-nlsRFP (red, nuclear signal). This section crosses the cell layer in a region of high-curvature. Only a band of two cell diameters can be seen in focus in this slice. The red and green stripe at the bottom left corresponds to the auto-fluorescence of the cuticle layer. The white line is used to generate the sagital section in c. **c.** Illustration of the *LocalZProjector* method. A 2D filter applied on each Z-plane is configured to generate a strong response at the layer of interest. Here the filter is a standard-deviation filter of window size 21×21. For each (X, Y) position, the Z-plane at which this filter has the strongest response is used to build a reference surface (white segmented line) around which intensity will be collected, possibly with an offset and a Z-range. All scale bars are 10 μm.

*LocalZProjector* relies on a few parameters set by the user. The height-map is determined by applying a 2D filter on each plane of the 3D source image, chosen and configured to yield a strong response for the layer of interest (either mean or standard deviation filter). To speed-up computation and reduce the effect of pixel noise, each 2D plane can be first binned. Noise can be further reduced through filtering with a Gaussian kernel. The height-map is then regularized using a median filter with a large window and rescaled to the original width and height. It is then used to extract a projection from the 3D image. A fixed `offset` can be specified separately for each channel, and is used to collect intensity in planes above or below the reference surface. Several planes, specified by a last parameter Δ*z*, can be accumulated to generate a better projection, averaging the pixel values or taking the maximum value of these planes (Figure 2c, Additional File 1: Movie S1).

Once the 3D volume has been projected on a single 2D plane, many tools are available that can segment the individual cells in the tissue. Several of them offer an intuitive user interface, allowing for immediate usage and user interaction. For instance, EpiTools (20) is a toolbox with MATLAB and Icy (21) components built to study the dynamics of *Drosophila* imaginal discs. Its segmentation algorithm relies on region growing from seeds determined automatically and merged based on region areas. SEGGA (22) is a standalone application written with MATLAB proposed for the investigation of *Drosophila* embryo germband epithelium. Recognizing that a small number of mistakes in segmentation can have major negative impact on cell tracking accuracy for long time-lapses, the authors propose several approaches to increase the robustness of segmentation and have a near perfect tracking results. TissueAnalyzer (23) is a tissue segmentation tool, distributed along TissueMiner (24) and the combination of these two softwares offers a framework that let end-users implement their own analyses using the R software and custom commands. EPySeg, a Python software that relies on deep-learning for the segmentation was recently made available in (25).

But these tools operate on 2D images only, which implies that the epithelium is a flat plane and parallel to XY. When this is not the case, most morphological measurements made on the 2D segmentation results will be corrupted by geometrical distortions induced by the projection (Figure 3a). Indeed, almost all morphology metrics, such as area, eccentricity and orientation will be affected when they are measured on the 2D projection of a curved or angled surface. *DeProj* aims at correcting these artifacts. It is a MATLAB tool that combines segmentation results and height-maps or 3D meshes to correct morphology measurements made on the 2D projection. *DeProj* returns corrected metrics, as if they were measured on the reference surface in the original 3D image (Figure 3b).

**Fig. 3.**
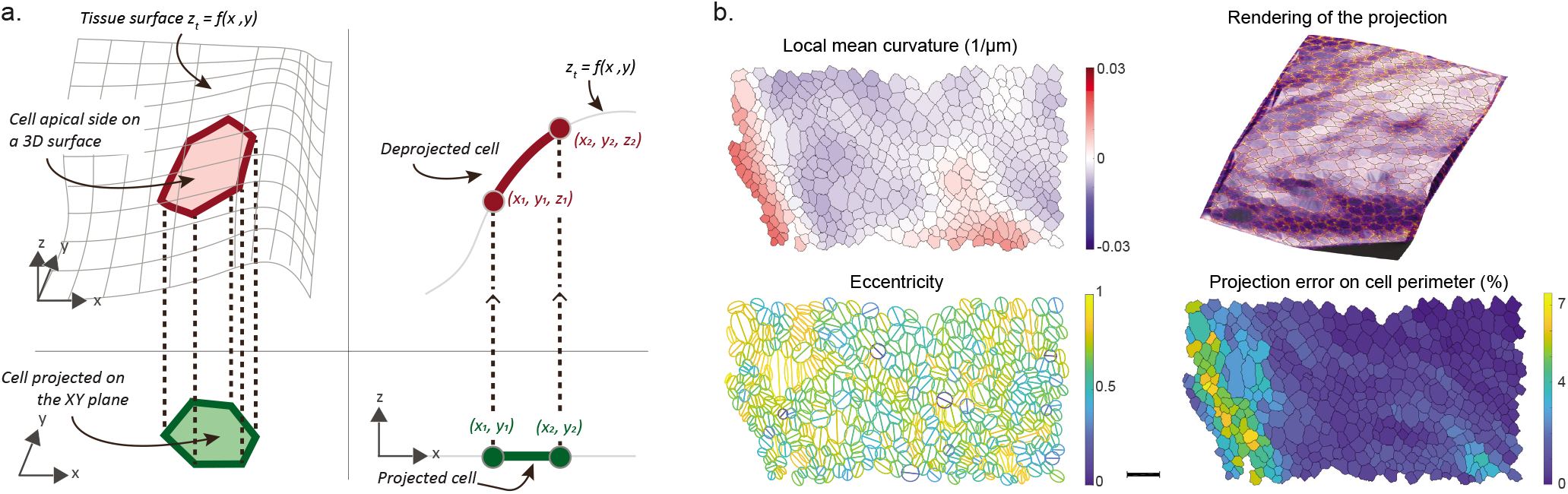
*DeProj*. **a.** *DeProj* is the second component of the toolbox and works by combining the tissue surface obtained *e.g*. via the height-map (gray line) and the segmentation of cells obtained on the 2D projection (green) to make measurements on the real cell contour in 3D (red). **b.** Example *DeProj* outputs, from left to right and top to bottom: the local mean curvature of the epithelium experienced by the cells; a 3D visualization of the epithelium 2D projection mapped on its 3D surface; the cell eccentricity measured in their oblique apical surface plane; the error (1 – *l*_2D_/*l*_3D_) comparing the cell perimeter measured on the 2D projection *versus* its real value inferred by *DeProj*. Scale bar: 10 μm.

## Results

### *LocalZProjector* is an accurate, fast and convenient tool to generate projections of 3D manifolds on 2D planes

One of our recent study involves long-term 3D timelapses imaging of a *Drosophila* pupal notum (3, 26). To follow the cells dynamics in the tissue, we relied on generating an accurate 2D projection of the cell monolayer (adherent junctions marked with E-Cadherin signal, Ecad-GFP), taken from the epithelium surface, and at the same time collecting the RFP signal from cell nuclei a few μm below this surface (nls-RFP). Imaging relevant signal in this tissue brings a few difficulties. First, the epithelium is not flat. Sampling a large portion of the tissue requires the acquisition of about 40-60 μm thick optically-sectioned volumes. Second, the cuticle (the exoskeleton of the fly) is located apically to the epithelium and is auto-fluorescent on a large spectrum (green to far-red). Third, below the epithelial layer lie large cells called fat bodies which are also highly fluorescent. In the GFP channel, the resulting 3D images display mainly three layers. The top one (low Z values) corresponds to the auto-fluorescent signal collected from the cuticle. The middle one corresponds to the epithelial cell layer, where the cell membranes build a manifold with a large curvature (Figure 2b). Below these two layers, the fat bodies generate punctate, bright structures. If not properly excluded from the projection, these unwanted structures in the imaged volume will degrade its quality and complicate or even forbid the subsequent segmentation task.

As the problem introduced here with the pupal notum is frequent in epithelium tissue imaging, we used it to validate the performance of the *LocalZProjector* plugin by comparing it to currently available projection tools (Table 1). We generated the user-expected 2D projection of the epithelium (ground truth) by manually selecting the junctional signal of epithelial cells in each plane (see Additional File 1: Supplemental Note 1 for details). We then calculated several metrics measuring the accuracy and performance of the projection generated by the *LocalZProjector* tool, and compared the results to 7 other methods (Additional File 1: Supplemental Note 2): the standard maximal-intensity-projection (*MIP*), the *StackFocuser* tool (9), *SurfCut* (12), *PreMosa* (10), the Extended-Depth-of-Field (*EDF*) tool (11), the Mininum-Cost-Z-Surface (*MinCostZ*) approach of (13, 14), implemented in (15) and the *Smooth Manifold Extraction* tool (8, 16). We first assessed how close the resulting projection was to the ground-truth projection using the root-mean square error (RMSE). This metrics proves useful in two aspects. First, because it allows for quantitatively comparing several methods and assessing how useful the projections will be in a subsequent analysis step. Second, because it offers a way to systematically search for optimal parameters for the 8 methods tested. For the comparison to be robust and fair, we ran indeed an extensive parameter sweep for each method, searching for the parameter set that gives the absolute best projection by minimizing the RMSE over a wide range of values (Additional File 1: Supplemental Note 2A). We then used this optimal projection for each method for the comparison. We find that the *LocalZProjector* projection is favored (Table 1, Additional File 1: Fig S3a and Additional File 1: Supplemental Note 2B) because it is robust against noise and can be configured to deal with regions of high-curvature. The accuracy of the height-map output is also important for the subsequent correction of the cell morphology measurements made by *DeProj*. We therefore calculated the RMSE of the height-maps compared with the ground-truth (Additional File 1: Supplemental Note 2C), for the projection tools that can return a height-map of the cell layer (*LocalZProjector*, *StackFocuser*, *PreMosa*, *EDF*, *MinCostZ* and *FastSME*). We find again that *LocalZProjector* offers the height-map with the lowest RMSE (Additional File 1: Fig S3b). Because *LocalZProjector* aims at being a tool possibly used on very long time-lapse movies, the time needed to generate a projection is important, we confirmed it is fast in comparison to most of the other methods (Additional File 1: Fig S3c). Finally, the projection accuracy of such a dataset is relevant mainly for its use in a subsequent analysis. We chose to focus on cell segmentation, as *DeProj* will be used to measure accurate and unbiased cell morphology. We therefore derived a simple, fully automated segmentation workflow on the projections, and compared segmentation results against a ground-truth segmentation (Additional File 1: Fig S3d, Additional File 1: Supplemental Note 3). We repeated the comparison presented in this table on a high quality image of a *Drosophila* pupa notum (Additional File 1: Table S1), and on an adult zebrafish brain image (Additional File 1: Table S2). The data, methodology and tools that support these comparisons are available online (27, 28). These results exemplify the usefulness of *LocalZProjector*, both for accuracy and performance.

**Table 1.** Features and performance of several end-users projection tools compared in this work. Most of the tools that are distributed within a framework like Fiji or MATLAB can be scripted or modified to harness the extra time and channel dimensions. This table reports whether they can do it without extra effort from the user. For the MIP technique we took the implementation in ImageJ. PreMosa has a separate command (ExtendedSurfaceExtraction) that can deal with multi-channel images. The 4 last columns relate the performance metrics of the tool measured with the *Drosophila* pupal notum image, and plotted in Additional File 1: Fig S3. Lower values indicate better performance. The *OCE segmentation* column reports the accuracy of the cell segmentation using object-consistency error metrics (see Additional File 1: Supplemental Note 3). The color scheme is determined from the range of results, splitting the range in 4 tiers, excluding the largest values for the height-map RMSE and timing metrics. The *MIP* does not return a height-map. On this image, *SurfCut* did not detect the epithelium, but the auto-fluorescent cuticle. By indicating a large shift in Z in the parameter, it could be made to return a usable projection nonetheless, but the height-map is aberrant and its RMSE measure is therefore not included in this table.

|  | References | Distribution | Reference surface detection | Multi-channel | Time-lapse | Large images | RMSE projection | RMSE height map | Timing (s) | OCE segmentation |
| --- | --- | --- | --- | --- | --- | --- | --- | --- | --- | --- |
| *MIP* |  | ImageJ plugin | ∅ | + | + | + | 304 |  | 0.06 | 0.538 |
| *StackFocuser* | (9) | ImageJ plugin | heuristics | - | - | - | 168 | 0.82 | 16.6 | 0.302 |
| *SurfCut* | (12) | ImageJ macro | heuristics | - | - | - | 129 |  | 6.0 | 0.187 |
| *PreMosa* | (10) | Standalone | heuristics | + | + | - | 144 | 0.51 | 4.3 | 0.208 |
| *EDF* | (11) | ImageJ plugin | heuristics | - | - | - | 236 | 7.24 | 127 | 0.405 |
| *MinCostZ* | (13–15) | ImageJ plugin | energy minimization | - | - | - | 114 | 1.25 | 6.2 | 0.198 |
| *FastSME* | (8, 16) | ImageJ plugin (SME),<br>MATLAB (FastSME) | energy minimization | + | - | - | 127 | 0.887 | 16.6 | 0.178 |
| *LocalZProjector* | This work | Fiji update site | heuristics | + | + | + | 81.5 | 0.428 | 4.3 | 0.117 |

Finally, in this study, we wanted to generate projections of the E-cadherin channel but also needed to visualise the nuclei reporter of some of these cells localised a few micrometres just below the reference surface, and follow it over time. This was easily feasible as *LocalZProjector* can handle multiple-channel long time-lapse images in a user-friendly manner. We compared some of the *LocalZProjector* capabilities with other tools in Table 1. We also proved qualitatively that *LocalZProjector* can be used on a wide-range of images coming from very different samples and we especially validated it on a large set of example images introduced in (16), taken from samples ranging from Neuroscience to Cell Biology and synthetic images. In Additional File 1: Fig S12 we present the projections obtained successfully with *LocalZProjector* on this dataset.

### Projecting large images with *LocalZProjector*

*In toto* imaging of developing embryos allows for investigating the dynamics of tissues at a large spatial scale. For instance when analyzing gastrulation in entire avian embryos, we showed that it is driven by the graded contraction of a large-scale supracellular actomyosin ring at the margin between the embryonic and extraembryonic territories (4). For this study we relied on particle image velocimetry (PIV) to measure the tissue displacement field. This technique does not require the segmentation and tracking of individual cells. However, several key mechanisms at large scales emerge from the dynamics of single cells (29). The need to segment all the cells in a whole embryo prompts for imaging at high resolution and special microscopes (30). But such acquisition setups generate in turn very large images. Also some imaging modalities that enable imaging large specimen at high resolution, such as Light-Sheet Fluorescence Microscopy (LSFM), may bring additional distortions in the image. In order to image a quail embryo at high resolution, we relied on LSFM using an inverted selective plane illumination microscope (30, 31).

While the light-sheet is held stationary at 45° of the embryo surface, the embryo is translated horizontally through the light-sheet. The 2D planes acquired for each translation are then concatenated in a 3D stack. Because the axis of the em-bryo translation and the light-sheet plane are at a 45° angle, the stack needs to be post-processed to remove the induced skew. The resulting is a 8669 ×2285 ×1067 image, amounting to a 42 GB file for a single time-point.

The 2D projection of such an image is very relevant as the epithelium of interest is a smooth and thin cell layer within a large 3D volume. It would also reduce the image size by 1067 and make it much more amenable to analysis and compact storage. To tackle the challenges coming with such large datasets, we use the Fiji *Virtual Stack importer*, which only loads one Z-slice in memory at a time. This allows for the opening and to some extent the processing of images much larger than RAM. While the MIP works well with virtual stacks, the resulting projection is corrupted by projection artifacts. Moreover, in such images the point-spread function (PSF) is not aligned with Z-axis of the image. The elongation of the PSF generates marked distortions and blurs the membrane signal in the projection, up to the point where cells cannot be outlined by eye (Figure 4a), even on a source image with little signal coming from spurious structures.

**Fig. 4.**
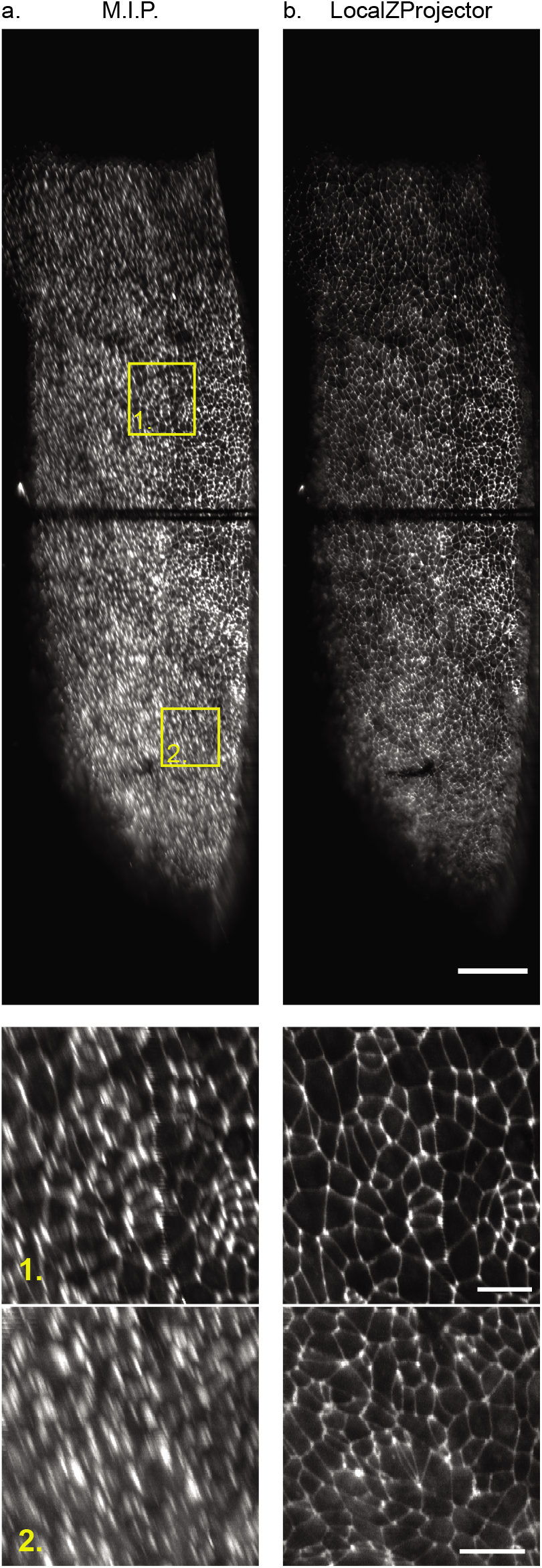
Projection of a large quail embryo (*Coturnix japonica*) imaged with LSFM. Projection results using the maximum intensity projection (**a.**) and the *LocaLZ-Projector* plugin (**b.**). Bottom: details of the two insets outlined in yellow in the top panels. Scale bars: top: 100 μm, bottom: 20μm. The two dark horizontal lines correspond to bleaching happening during the setup phase of the experiment. The intensity display range is the same on the 6 images.

To improve the projection quality, we developed the *LocalZProjector* so that it can work with virtual stacks as well, granting it the ability to project images larger than the available memory amount and without having to do any preprocessing or resaving of the image. *LocalZProjector* works with two passes through the image data, one to compute the reference surface, one to perform the local projection (Additional File 1: Movie S2), ensuring that individual planes are read from disk at most twice. This still comes at a time penalty. *LocaLZprojector* takes 8.5 minutes per time-point, against 1.3 minutes for the MIP. But the *LocalZProjector* result is completely devoid of the defects observed with the MIP and is amenable to segmentation and quantification (Figure 4b).

### Accurate measurements of cell morphology: *DeProj*

In vertebrates, adult neural stem cells (NSCs) are responsible for adult neurogenesis (32) and, in some vertebrates, regeneration post-injury (33). NSCs are organized as an epithelial-like structure lining ventricles that has to be maintained functional for very long periods of time (often over years). To understand NSC population homeostasis, it is essential to integrate large-scale and long-term imaging of the NSC pool and the zebrafish telencephalon has recently emerged as a unique model for these studies (34, 35). To study cellular and mechanical functions of NSCs over the entire dorsal telencephalon (pallium) we can image whole-mount immunostainings against ZO1 (a component of tight junctions (5)), highlighting apical domains of the NSCs. However, since the pallial hemispheres are highly curved we so far could not extract the geometrical parameters of many NSCs (falling in periphery, in sulci, *etc*.).

*DeProj* is a MATLAB app specifically built to address this issue. *DeProj* requires the reference surface that gives the shape of the tissue in the form *z_t_* = *f*(*x*,*y*). The reference surface can be specified as the height-map which is the secondary output of *LocaLZProjector* and several others projection tools, or as a mesh that extends over the epithelium surface. It is smoothed by a Gaussian filter with a *σ* equals to the median diameter of the cells, to mitigate step-wise patterns, happening for instance when the height-map is made of integer Z-slice positions. *DeProj* then takes the 2D output of the segmentation results, and creates cell objects as closed polygons. For each point of the polygon, a *z* position is determined from its (*x*,*y*) coordinates from the reference surface function. A new 3D polygon is then created, that represents the cell apical surface in 3D, effectively deprojecting it on the tissue surface. The cells segmentation can be specified as a black and white mask, or as a MATLAB structure returned *e.g*. by TissueMiner (24) or the tool of (36), and documented online. Several morphological metrics (area, perimeter, orientation, eccentricity, number of neighbors, fit by an ellipse in 3D, Euler angles, local curvatures) are then computed and saved, along with the cell contour mapped on the tissue surface. The generated *DeProj* data object is used to store the analysis results and offer exporting facilities and several visualisations of the results.

On the telecephalon image, a 3D view generated by *DeProj* shows the shape of the epithelium. We can see several regions where the tissue is very curved, particularly at its borders and in the sulcus separating the regions (Figure 5a). A visual representation of the cell area does not show a salient difference between a measurement made on the proper 3D epithelium surface (Figure 5b) or on the 2D projection (Figure 5c). However the histogram of the error metric *e_a_* = (1 – *a*_2D_/*a*_3D_) between these two quantities shows that for a large number of cells, using the 2D measurement induces an error greater than 20% (Figure 5d, 22% of the 3000 cells in this epithelium have an error larger than 20%). The cells with a large error are found at the epithelium border and in the sulcus (Figure 5e), which are regions where the angle between the cell apical planes and the XY plane is especially large (Figure 5f). Without surprise, we find that a large slope correlates with a large error (Figure 5g). If a cell would be a square of side *a*, with one side making an angle *θ* with the XY plane, then its real area measured in 3D is *a*^2^. The 2D projection of this cell contour on the XY plane generates a rectangle of sides *a* and *a* × cos*θ*, so that the error *e_a_* is equal to 1 – cos*θ* for this cell. Because real cells have complex shapes and have a contour that is not necessarily contained in a plane, we find that this expression constitutes a lower bound for *e_a_* (Figure 5g, red line).

**Fig. 5.**
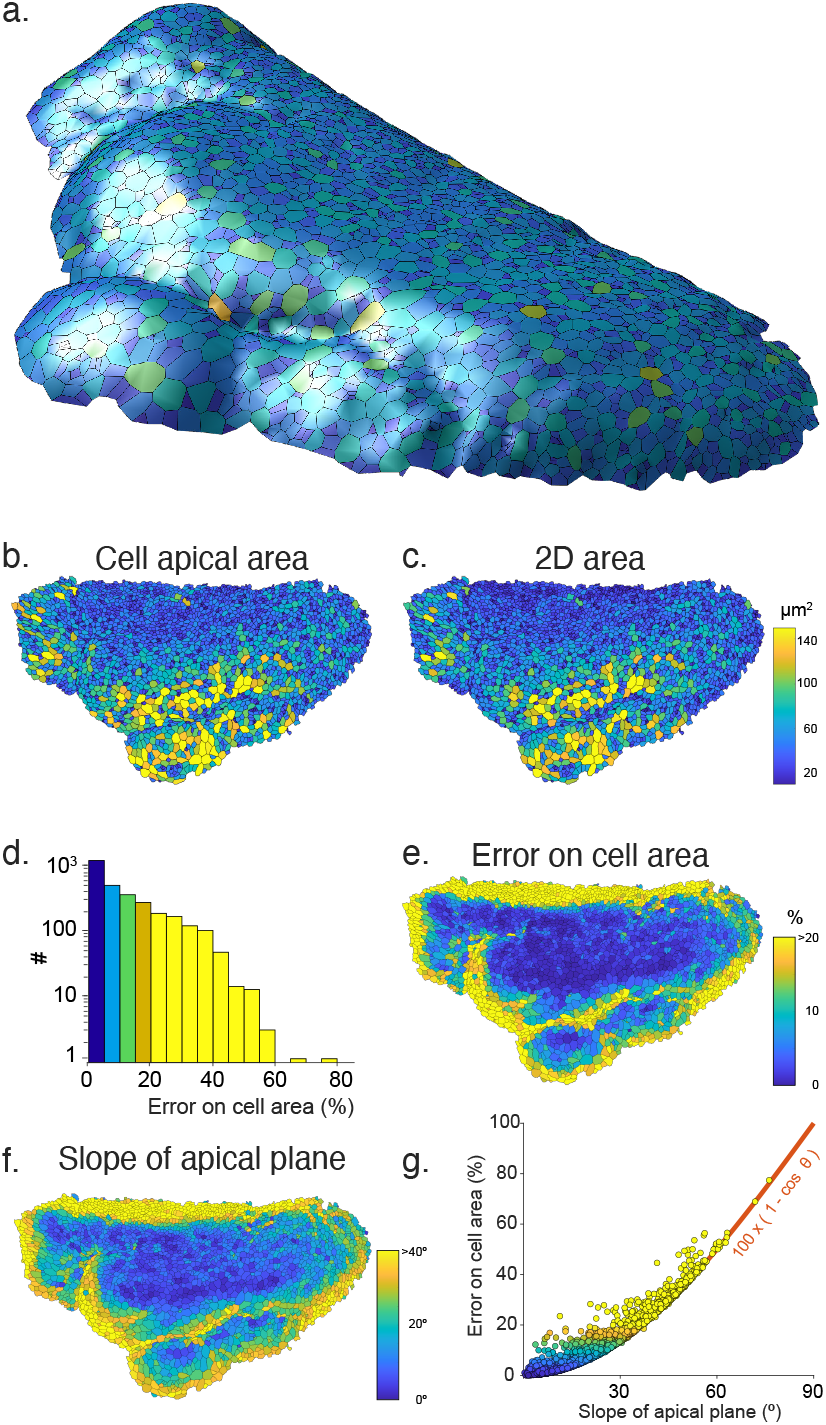
Getting accurate cell morphology measurements in non-flat samples with *DeProj*. **a.** 3D visualization of the NSC population on the zebrafish telencephalon generated by *DeProj*. The cells are drawn with their approximate contour, their color encoding the number of neighbor cells (from dark blue to yellow: 2 to 16 neighbors). **b.** Cell apical area, measured on the 3D surface, *a*_3D_. **c.** Cell apical area measured on the 2D projection, *a*_2D_. The color scale is identical in b and c. **d.** Histogram of the error metric on area (1 – *a*_2D_/*a*_3D_) for all the cells of the epithelial-like surface. **e.** Rendering of this error on the epithelial-like surface. **f.** Rendering of the slope of the apical plane of each cell with the XY plane. **g.** Correlation between the slope of the apical plane and the error on cell area for all cells of the epithelial-like surface. Red line: 100 × (1 – cos*θ*).

The correction offered by *DeProj* is only accurate if the height-map or the tissue mesh follow the tissue surface accurately. We have seen above that projection methods can return height-maps that deviate from the ground-truth (Table 1). Of course, a uniform error in the Z positions returned by the height-map won’t impact the cells morphology measurements. Similarly, an error that is uniform over a spatial scale larger than a cell will have only a limited effect. We must therefore consider a difficult scenario, where individual cells are deformed by an erroneous height-map. To assess the impact of such a situation, we took the data of Figure 5 and artificially moved the Z position of half the points along the boundary of each cell by one Z-slice, keeping the other half in place. We then compared the 3D area and 3D perimeter measurements made with and without altering the cell contour. We found that the error induced is 4.2 ±3.5 % on the area measurements and 2.4 ±2.0 % on the perimeter. This error range is smaller than what we aim at correcting for with *DeProj* (Figure 5d), but is important enough to recommend checking for the accuracy of the height-maps with the same scrutiny than for the projections.

## Discussion

As can be noted in the comparative study of this work, there exists already several tools that perform projection of tissues in 2D from a 3D image. Their number demonstrates the importance of the information that can be extracted from the resulting images. This, and the still popular usage of MIP despite its shortcomings, also points out the difficulty of having a tool that can address all types and qualities of images to project, despite the similarity in tissues staining and shape. Yet the quality of projection dictates the subsequent step in analysis, as demonstrated in Additional File 1: Fig S3d. Some of these tools have the advantage of being parameter-free (8, 13–16). *LocalZProjector* takes another approach and requires several parameters to be tuned. In turn, this configuration step allows it to work even with difficult 3D images containing spurious structures and confers it a greater adaptability. It is also to our knowledge the only one that can process large images without pre-processing.

*DeProj* allows for the correction of geometrical distortions caused by the projection on morphological measurements. These artifacts remain often overlooked, despite possibly compromising the measurements accuracy when the local angle of the tissue with the XY plane is large. It works as the final step in our bioimage analysis pipeline and combine the cell segmentation results with the original shape of the sample, such as a height map, to yield various corrected tissue visualizations and accurate morphological measurements on cells. Typically microscopists prepare samples in such a way that the orientation of the tissue is favorable for imaging, with its main orientation parallel to the XY plane. Yet, we found that in a tissue like the pallium, the slope can exceed 40° in the regions of interest. But even a more moderate slope yields dramatic errors on measurements taken directly on the 2D projection (Figure 5g). *DeProj* offers robustness against these distortions, and makes it possible to accurately access the morphology of cells in highly curved samples while taking advantage of a simplified 2D dimensionality to segment the tissue morphology.

## Conclusions

The flexibility of *LocalZProjector* enables obtaining high-quality 2D projections even when the source image display spurious structures, a high curvature, low signal-to-noise ratio, and is of large size. High quality projections are required for the robustness of subsequent analysis such as cell segmentation. *DeProj* is an ideal companion tool in a cell segmentation pipeline, as it allows for retrieving accurate morphology measurements, even on the 2D projections of highly-curved samples. *LocalZProjector* and *DeProj* constitute together a useful toolbox to get accurate cell morphology measurements on epithelial tissues.

## Methods

### Drosophila imaging

Notum live imaging was performed as described previously (3). Briefly, the pupae were collected at the early stage (0-6 hours after pupal formation), aged at 29°C, glued on double sided tape on a slide and surrounded by two metal spacers of approx. 0.650 mm. The pupal case was opened up to the abdomen using forceps and mounted with a 20 × 40 mm ##1.5 coverslip where we buttered halocarbon oil 10S. The coverslip was then tapped on the spacers using regular tape. Pupae were collected 48 or 72 h after clone induction and dissected 16-18h after pupae formation (APF). Pupae were imaged at 29°C for 22 hours on a LSM 880 scanning laser confocal microscope (Carl Zeiss A.G.) equipped with a fast Airyscan module using an oil 40X objective (NA 1.3), Z stacks (1 μm/slice), every 5 min using autofocus. The autofocus was performed using the autofluorescence of the cuticle in far red (using a Zen Macro developed by Jan Ellenberg laboratory, MyPic). Movies were performed in the nota close to the scutellum region containing the midline and the aDC and pDC macrochaetae. The experiment presents a pupae with endoCad::GFP signal and groups of cell overexpressing UAS-yorkie S11A S168A S250A V5 clones and nuclei in red over the control of the GAL80TS thermosensitive. The cross and the progeny were kept at 18°C, and the pupae were switched to 29°C 8 hours prior to the movie for conditional activation.

### Quail embryo imaging

Fertilized quail eggs (Coturnix japonica) were purchased from Cailles de Chanteloup. The embryos were collected at stage XI, fixed in 4% formaldehyde/PBS for 3 hours at 4°C and washed / blocked in PBS / 0.1% Triton X-100 / 2% BSA (from Roche)/10% FBS (from Gibco). The primary antibody used was ZO-1 (Invitrogen ZO1-1A12) at 1:200 and the secondary antibody was goat anti-mouse AlexaFluor 488 (A28175) at 1:500. The embryos were mounted with DAPI-containing Fluoromount-GTM (eBioscience) between slide and coverslip and sealed with commercial nail polish.

Fixed quail embryos were imaged using a Dual Inverted Single Plane Imaging Microscope (DiSPIM, 3i Marianas Light sheet). The system geometry consists in two identical arms containing an illumination path composed of laser light output directed to a XY scanner for generating the light sheet and a detection path where an sCMOS camera (Hamamatsu Orca Flash 4) is fitted. Both arms are assembled at 90 degrees and alternate from stimulating the embryo to detecting its opposite side. This ensemble is rotated by 45 degrees which allows to work on flat mounted sample (31). Those arms are fitted with 40x 0.8NA water immersion objective (Nikon CFI Apo NIR 40x W) with a 3.5 mm working distance. The field of view of the objective is 330 mm and the depth detection is limited to scattering and absorption of the light within the sample (around 30 μm in our sample). The embryo was irradiated with 561 nm excitation light and the stage was scanned through both light-sheets simultaneously across 1.408 mm with a step size of 0.42 mm. The resulting acquired stacks dimensions are 1024 x 2048 x 3339 pixels. The image stack produced by this system was then de-skewed and rotated by 45° to yield a proper geometrical representation of the sample. After de-skewing and rotation, the final image dimension are 8669 × 2285 × 1067 pixels.

### Adult zebrafish brain imaging

Brains were dissected in 1X solution of phosphate buffered saline (PBS - Fisher Bioreagents) and directly transferred to a 4% paraformaldehyde solution in PBS for fixation. They were fixed for 2 to 4 hours at room temperature (RT) under permanent agitation. After four washing steps in PBS, brains were dehydrated through 5 minutes series of 25%, 50% and 75% methanol diluted in 0.1% tween-20 (Sigma Life Science – P9416) PBS solution and kept in 100% methanol (Sigma-Aldrich, 322415) at −20°C. The whole-mount immunohistochemistry (IHC) started by the rehydration of the telencephali. Then, the brains were subjected to an antigen retrieval step using Histo-VT One (Nacalai Tesque) for an hour at 65°C. Brains were rinsed in a 0.1% DMSO and 0.1% Triton X-100 (Sigma Life Science– 1002135493) PBS 1X solution (PBT) and then blocked with 4% normal goat serum in PBT (blocking buffer) 4 hours at RT. The blocking buffer was later replaced by the primary antibodies solution, and the brains were kept overnight at 4°C on a rocking platform. The next day, brains were rinsed over 24 hours at room temperature with PBT and incubated in a solution of secondary antibodies diluted in PBT overnight, in the dark, and at 4°C on a rocking platform. After several washes, the telencephali were mounted in PBS on slides using a 0.7 mm-thick holder. The primary antibody anti-ZO1 was used at 1:200 (Mouse monoclonal IgG1 anti-ZO1, Thermo Fisher, cat. #33-9100, RRID: AB_2533147) and the secondary antibody anti IgG was used at 1:1000 (Goat anti-Mouse IgG (H+L) Alexa633 conjugated, Thermo Fisher, cat. #A-21052, RRID: AB_2535719).

Images of whole-mounted immunostained telencephali were acquired on confocal microscope (LSM700, Carl Zeiss A.G), using a 40X oil objective. We acquired images with a z-step of 0.65 μm. We averaged each line four times with an image resolution of 1024 ×1024 pixels with a bit-depth of 12-bits. The power of the lasers was kept constant for all of the acquisitions and the gain was adjusted for each experiment. We recorded mosaics with a 15% overlap to image an entire hemisphere per fish.

## Declarations

## Acknowledgements

We thank Stéphane Rigaud for his help with the software from (36).

## Consent for Publication

Not applicable.

## Availability of Data and Materials

The project homepages contain the source code, installation instructions, documentations and extra implementation details. For *DeProj*, the homepage also contains 3 example scripts and data to reproduce some panels in Figures 1, 2 and 5. Part of the raw data of this study and several examples are available on Zenodo (27, 37, 38). The tools and methodology to determine optimal parameter sets and performance metrics for the 8 methods compared in Table 1 are available and documented, along with an example dataset, on Zenodo as well (28). The rest of the data is available upon reasonable request.

### Local Z Projector

- *Project name*: LocalZProjector.
- *Project homepage*: https://gitlab.pasteur.fr/iah-public/localzprojector
- *Operating systems*: Platform independent.
- *Programming language*: Java.
- *Other requirements*: Runs from Fiji (19).
- *License*: BSD 3
- *Any restrictions to use by non-academics*: None.

### DeProj

- *Project name*: DeProj.
- *Project homepage*: https://gitlab.pasteur.fr/iah-public/DeProj
- *Operating systems*: Platform independent.
- *Programming language*: MATLAB.
- *Other requirements*: at least MATLAB R2019b and the Image Processing Toolbox.
- *License*: BSD 3
- *Any restrictions to use by non-academics*: None.

## Authors’ contributions

SH, LV and JYT wrote the code. SH, LV, LM, ND, PC, EE and JYT performed the experiments. SH, LV and JYT wrote the article with contributions from all authors. All authors read and approved the final manuscript.

## Ethics approval and consent to participate

Not applicable.

## Additional Files

**Additional File 1: Supplemental Notes 1-3, Supplemental Figures 1-12.**

***Supplemental Note 1.*** Generation of a ground-truth dataset.

***Supplemental Note 2.*** Comparing projections against the groundtruth dataset for several methods.

***Supplemental Note 3.*** Comparing segmentation results on optimal projections.

***Table S1.*** Comparison of projection performance metrics on a high quality image of a *Drosophila* pupa notum.

***Table S2.*** Comparison of projection performance metrics on the adult zebrafish brain image also used in Figure 5.

***Figure S1.*** Overview of the Local Z Projector method.

***Figure S2.*** Manual generation of a ground truth dataset.

***Figure S3.*** Performance metrics comparison for projection tools.

***Figure S4.*** Optimization of the parameters of the Local Z Projector method.

***Figure S5.*** Optimization of the filter-size parameter of the Stack Focuser method.

***Figure S6.*** Optimization for the parameters of the PreMosa method.

***Figure S7.*** Optimization for the parameters of the Extended Depth of Field method.

***Figure S8.*** Optimization for the parameters of the Minimum Cost Z Surface method.

***Figure S9.*** Optimization for the parameters of the FastSME method.

***Figure S10.*** Projection results comparison for 8 methods.

***Figure S11.*** Height-map results comparison for 6 methods.

***Figure S12.*** Qualitative performance of LocalZProjector on other datasets.

**Additional File 2: Movie S1.** Capture of a local projection process of the drosophila pupal notum. Top, from left top right: Individual Z-slices of the 3D stack of the Cadherin-GFP channel. Mask indicating what part of the current Z-slice belongs to the reference surface. Part of the current Z-slice within the mask. Bottom, from left to right: Resulting local projection of the Cadherin-GFP channel (left), of the miniCic ERK biosensor (middle, Scarlet fluorescent protein), and of a nuclear far red protein (right, iRFP1.0 fluorescent protein) centered 3 μm below the reference plane (for the middle and right projections).

**Additional File 3: Movie S2.** Capture of the local projection process for the quail embryo image shown in Figure 4b.

