## Supplemental information for "*LocalZProjector* and *DeProj*: A toolbox for local 2D projection and accurate morphometrics of large 3D microscopy images"

### Supplementary Note 1: Generation of a ground-truth dataset.

A ground-truth dataset is required to compare the *LocalZProjector* tool results with other methods. To generate this ground truth, we selected a single time-point in the *Drosophila* pupal notum dataset of the main text. This single-channel 3D stack displays two salient layers. The top one (small Z values) is generated by the autofluorescence of the cuticle. It is noisy, continuous and shows up in all fluorescence channel. Below the cuticle, we find the cell layer, where the cells apical junctions are labeled for their membrane by Ecad::GFP. The cuticle layer and the cell layer are separated by about 4 to 9 slices. Under the cell layer (largest Z values) there are about 10 locations where discrete and bright structures can be seen, corresponding to apoptotic cells. In the right part of the image, the cell layer is almost parallel to the XY plane and is located just under the cuticle layer. The left part of the cell layer displays a relatively high curvature (Figure 1b).

We generated the ground-truth dataset using Icy [1]. Each slice of the 3D stack was manually annotated for regions where the cells of interest are in focus (Figure S2). A Jython script then parsed these regions to fetch the pixel in the specified Z-plane for each (X,Y) position, and writes its value on a single plane, generating the ground-truth for the projection. When a single (X,Y) position was represented by several regions in different Z planes, the pixel with the maximal value was retained. The height-map image corresponding to this projection was generated along, by simply writing the Z position retained for projection as pixel value. When a (X,Y) position was represented by several regions in different Z planes, the height-map value was taken as the mean of all the Z "in-focus" positions. This led to the height-map ground-truth image having non-integer values for some pixels. The resulting projection is a 2D gray-scale image of size 1148 ×1148. Its bottom-left corner is devoid of signal and was excluded from all the metrics described in this work.

### Supplementary Note 2: Comparing projections against the ground-truth dataset for several methods.

The performance of 8 different projection tools were then compared using the *Drosophila* pupal notum image and the ground-truth image generated as detailed above. We ran an extended parameter search, in order to determine the parameter value that give the best projection for each method. This provides valuable know-how as how a user can select values based on the input image. We put this information in the documentation of *LocalZProjector*, on its homepage. We give in this section the details of the performance comparison. The results are summarized in the Figure S3 and in the Table 1.

**A. Optimizing projection parameters for the compared methods.** We want to assess how a projection method can be useful in practice for end-users, in retrieving a 2D projection that can be used in subsequent analysis steps. Therefore, we first turn to a pixel-based comparison of the 2D projection given by each method with the projection ground-truth. We use the RMSE to report these differences, defined here as:

$$\text{RMSE}_M = \sqrt{\frac{1}{N_{\text{pixels}}} \sum_i \left(p_M(i) - p_{GT}(i)\right)^2} \tag{1}$$

where $N_{\text{pixels}}$ is the number of pixels in the projected image, $p_M(i)$ the pixel value in the projected image by the method $M$, $p_{GT}(i)$ the pixel value in the projection ground-truth and $i$ iterating over all the pixels in each image. We excluded from the RMSE calculation the area of the sample where there is no cell layer.

Several of the methods tested allow to define a certain thickness parameters specifying how many Z-planes around the Z position defined by the height-map to include in a local projection. This is the case for the *LocalZProjector* and for *Min Cost Z Surface*. This offers some tolerance against small inaccuracies of the height-map. When generating the ground-truth projection, we take the brightest pixels of several Z-planes when several of them are annotated as being in focus. Because of this, methods that can accumulate several planes are slightly advantaged.

Moreover, the use of manually annotated ground-truth dataset depicted above as a basis for performance comparisons has some limitations. Indeed, the metric values we get might be affected by the imprecision of manual annotation. We nonetheless use the RMSE defined above to rank each method and detect projection defects, keeping in mind the aforementioned limitation. We also can use the RMSE metric to optimize the parameters of each method separately and derive practical advice on how to tune them with respect to the image content.

***Maximum intensity projection.*** We used the *Z Project...* command of ImageJ [2] in the Fiji distribution [3], choosing maximum intensity as a projection method. It generates a projection by taking the pixel with the largest value along a (X, Y) column. This method is the simplest projection technique and does not have parameters to tune.

***Local Z Projector.*** The *LocalZProjector* aims at being a general tool and has therefore several parameters that can be tuned to accommodate a large variety of input. We derived the optimal set of values for these parameters by parameter sweeps, testing a large number of parameter combination (close to 200,000) and retaining the combination yielding the lowest RMSE. For all the tests described below we used the `Max of Std` method to generate the height-map, and the `MIP` method to accumulate several planes around it. We have found that the lowest RMSE value is 81.5 and obtained for binning = 2, gaussian sigma = 0.5 pixels, filter size = 13 pixels, median filter half-size = 16 pixels and $\Delta z$ = 1. The projection takes about 5 s. to complete with these parameters. A large subset of parameters give also results close to the optimum, but in a much shorter time. For instance the parameters binning = 5, gaussian sigma = 0 pixels, filter size = 7 pixels, median filter half-size = 4 pixels and $\Delta z$ = 1 yields a RMSE of 83.0 in about 0.6 s. Nonetheless, we base the following discussion on parameter values associated to the absolute optimum.

The binning value has a dramatic positive impact on processing time (Figure S4a). A large binning can be chosen without compromising the RMSE value significantly. For very large binning, the structures that define the layer of interest (in our case the manifold shape of the cell membranes) are confused in the downsized image and cannot be retrieved later by the standard deviation filter.

The Gaussian filter is meant to limit the effect of noise. But binning already averages pixels together when downsizing. Even with a binning of 2, the Gaussian filter has almost no impact on RMSE or timing. (Figure S4b).

The main filter is the key parameter to tune to reliably retrieve the cell layer in the projection and not the cuticle layer. Its type must be chosen to generate a strong response for the cell labelling. In our case the layer we are interested in displays cells as manifolds, and the cuticle generates a spurious layer where the signal is very slowly varying (minus the noise). An adequate filter is therefore the standard-deviation filter, that will act as a measure of a local contrast between the bright cell membranes and the dark cell centers. Its size must be chosen so that the contrasted features will be fully included in one window. The cells sections in the test dataset are about 20-40 pixels in diameter, with a membrane thickness around 3-5 pixels. We find that the optimal value of the size is a value of 13 pixels, enough to include all or a large part of a single cell (Figure S4c). Smaller values would create too many windows without contrast. For larger values, the cuticle layer gives a strong response in the areas of the sample where the curvature is large. The projection then displays jumps between the two layers. In the *LocalZProjector* plugin, the size of the main filter is specified independently of the binning value. It is then scaled to match the size in the downsized image. This is why the RMSE plot for this parameter in the Figure S4c displays a staircase pattern with steps corresponding to the binning value.

A median filter can be applied as a post-processing option on the height-map image. It is meant to remove spurious Z value that can be generated by punctate structures above or below the layer of interest. Its size has to be therefore large compared to these structures, and small compared to the layer curvature. In our main example, such bright fluorescent structures appear from some dead cell bodies away from the epithelial surface, and are picked by the standard-deviation filter, generating locally spurious large Z values. A median filter of half-size of 16 pixels in the downsized height-map completely remove these spurious jumps (Figure S4d). This corresponds to a size in the original scale image of 64 pixels, about twice the cells diameter.

Finally, the $\Delta z$ parameter specifies how many planes to accumulate around the reference Z plane for each (X, Y) position.

This allows for accommodating small mistakes in the height-map. In our case a value of $\Delta z \pm 1$ has a strong positive impact, dividing the RMSE by a factor of about 2 compared to no accumulation. However large values of $\Delta z$ will result in including unwanted signal from planes that are away from the reference surface. Unsurprisingly we find that a value of $\Delta z = 1$ returns the best projection (Figure S4e).

***Stack focuser.*** *Stack Focuser* [4] is an ImageJ plugin that generates projections by selecting the best plane for each (X, Y) column. This best plane is determined by applying the Sobel filter [5] at each Z-plane and selecting the one for which the filtered value is the largest. The size of the Sobel filter can be adjusted, and we found the optimal value to be 25, in the range of the cells size (Figure S5). For smaller values, the projection includes signal from the cuticle inside the cells and resembles that of *MIP* method. For large values, strong defects appear in the regions of the cell layer where the curvature is large.

***SurfCut.*** ImageJ *SurfCut* [6] is an ImageJ macro that extracts a projection by thresholding single Z-planes from a Z-stack, then stacking the obtained binary masks to generate a 3D binary mask of the sample. The top surface of this mask is then used to define a reference surface. With this approach SurfCut detects the cuticle layer and not the cell tissue layer. However the SurfCut tool allows defining offsets from the reference surface to generate the projection. Since the cuticle layer is almost parallel to the tissue layer, we could find parameters that would still generate projections of the tissue layer itself. Because SurfCut only operates on 8-bit images, we converted the source image to 8-bit, then re-scaled the projection so that the RMSE values reported in the Figure S3a is commensurate with values from other methods. We used the following parameters to do so: filtering radius = 6, threshold = 10, offset top = 5, offset bottom = 12.

***PreMosa.*** *PreMosa* [7] is a standalone command-line tool written in C++ that was initially developed to facilitate the pre-processing of large mosaics of the *Drosophila melanogaster* developing wing. It offers a tool to generate a projection from 3D stacks, a tool to stitch a collection of 2D projections into a large mosaic, a tool to correct uneven illumination and a tool to adjust the contrast. Since our sample is a single Field of View, we only tested and compared the former.

The surface projection tool of *PreMosa* works by first sub-dividing the image in square columns of size $r \times r$, $r$ the grid size being an adjustable parameter. For each column, a candidate Z-position is obtained by taking the plane in which there is the largest amount of bright-pixels. Bright-pixels are pixels that have a value larger than the lowest intensity of the top $\lambda$% pixels in the column, 20% by default. The bright-pixel threshold $\lambda$ is a second adjustable parameter. The resulting surface is then iterably smoothed and refined by determining the Z-position of greatest contrast (local variance), in a neighborhood of $\pm d$ planes around the surface. The maximal distance $d$ is a third adjustable parameter.

The grid-size parameter appears to be the crucial parameter to tune (Figure S6). For too small values of the grid-size, the projection is principally made of the cuticle layer, with small regions containing the cell layer. For values too large, the projection fails to include the curved regions of the cell layer. The distance parameter $d$ and the threshold $\lambda$ haves only little influence on RMSE for reasonable values of $r$.

***Extended Depth Of Field.*** The *Extended Depth Of Field* method aims at reconstructing a projection from a 3D stack by merging areas that are detected to be in-focus as the Z-position varies [8]. This algorithm is made to excel for transmitted light images (*e.g.* bright-field images or colored images from histology samples) and reflected, wide-field images. It is not made to deal with optically-sectioned, confocal stacks of structures labelled in fluorescence, where the signal coming from a structure is only present in a few Z-slices around its position. However the tissue layer in our test dataset contains cells labelled for their membrane. For the Z-planes that contain them, they will appear as sharp and well contrasted structures, and the in-focus detection of this algorithm may select them correctly.

The authors of [8] has made the technique available *via* an ImageJ plugin that we used in our comparisons. It offers a very large range of parameters that can be tuned, from the various techniques to detect in-focus plane and the parameters that configure them, to the topology smoothing steps. We limited our parameter sweeps to parse all the possibilities offered by the simple version of the user-interface, namely 5 possible values in the `Quality` settings and 5 possible values in the

`Topology` settings (see Figure S7). We note that for our tests, the topology parameter set does not have an impact on the output. Indeed, the range of smoothing or regularization associated with these parameter sets is relatively small compared to the size of our test dataset features. Accordingly, the height-map returned by any of the parameter set tested is very irregular. The `Complex wavelets with MCC` quality parameter set however gives a good response to the membranes of the cell layer. But indifferently of the topology parameter we use, the centers of the cells always contain spurious signal in the final projection.

***Min Cost Z Surface.*** The 5 methods above are direct methods that determine a best Z-plane for every (X, Y) pixel based on sharpness, brightness or local contrast. Some of them offer post-processing steps to smooth or regularize the resulting reference surface. Another approach consists in formulating the problem as a global energy-minimization. In [9] and [10], the authors use graph-cuts to extract a single smooth surface [9] or a pair of smooth surfaces that are roughly parallel [10] and made of bright pixels. The latter approach should be well suited to deal with our test dataset, as it displays two almost parallel layers.

The algorithms described in [9] and [10] have been implemented in Fiji in a plugin called *Min Cost Z Surface* [11]. Our attempts with the single-surface extraction method only resulted in a projection made of patches taken from both the cuticle layer and the cell layer with large gaps and jumps in the height-map. Changing the downsampling from x2 to x4 changes the patches that are incorporated but does not fix these defects. However, using the two-surfaces detection and with some tuning of the input parameters (maximal separation of the two surfaces, number of planes to accumulate in the final projection) we extracted separately the cuticle layer and the cell layer rather reliably. The optimum is obtained by incorporating 3 slices around the reference surface. The optimal value for this last parameter is found to be the same than for the $\Delta z$ parameter of the *Local Z Projector* method (Figure S8).

***Fast SME.*** The Smooth-Manifold-Extraction algorithms offers a projection technique that is also based on a global energy-minimization approach and has the advantage of being parameter-free [12, 13]. It can harness both 3D confocal stacks and 3D wide-field stacks. It is especially developed to deal with single cell monolayer labeled for their membrane. It is therefore one of the best suited tools to our test dataset. Briefly, the algorithm relies on classifying (X, Y) columns as belonging to the foreground (they contain a pixel belonging to the cell membrane) or belonging to the background (inside cells). The reference surface is then obtained energy minimization. The energy to minimize is the sum of a local smoothness term ensuring the regularity of the surface, and a distance term driving the surface close to the Z-planes of the foreground columns, and that ignores the background columns. A first paper [12] contains the original implementation, which was further optimized for better performance [13]. We base our comparisons on the latter, which is implemented in MATLAB. The user must only specify whether the image comes from a confocal imaging system (our case) or from a wide-field system, and how many slices to incorporate in a local projection around the reference surface. To emulate what we have been testing for the *Local Z Projector* and *Min Cost Z Surface method*, we varied this last parameter to include just the plane from the reference surface, $\pm1$ and $\pm2$ planes (Figure S9).

**B. Comparing optimal projections.** We then compared the optimal projection of each method against one another. The relative RMSE values are reported in the Figure S3b and the Table 1 in the main text. In the Figure S10 we report the 2D optimal projection for each method, along with the error map. The error-map is displayed as the pixel by pixel absolute difference with the value in the ground-truth projection, filtered by a $41 \times 41$ median. Except for *MIP* and *Fast SME*, all of the presented methods have some degree of parameter configuration that we could optimize to get a projection close to the ground-truth. Therefore all the optimal projections are satisfactory at least in some part of the image.

The *MIP* method incorporate the brightest pixel in the projection regardless of its plane of origin. Because of this, the signal inside the cells in the projection is made of bright pixels coming from the pupal cuticle signal, which is noisy and bright. Consequently the error map image shows large errors everywhere in the projection.

The *LocalZProjector* could retrieve a projection that is very close to the ground-truth, except in the flat region of the tissue (right arrow). In the latter region, the cuticle layer just above the tissue layer emits a strong signal. It is likely that here, projection includes spurious signal from the cuticle. Indeed, the parameters we picked as such that the local projection takes the maximum of 3 pixels around the reference surface.

The *Stack Focuser* plugin bears some similarities with the *LocalZProjector*. It determines the position of the reference surface by a filter that has a strong response to local contrast. Since in our case the cells membrane amount to salient ridges, this method correctly picks the cell layer. However, it does not offer a step that would regularize the reference surface, and the projection is deformed by some bright and punctate structures under the cell surface (arrows). Finally, in regions where the curvature of the tissue layer is large, the Sobel filter does not pick the right planes.

As mentioned above, *SurfCut* actually picks the cuticle layer. We configured it so that projection includes the pixels below the cuticle layer, hereby including those of the cell layer. Because the reference surface follows the cuticle layer and not the cell layer, areas of the projection where the error is large actually reflects irregularities in the former.

*PreMosa* detects candidate planes by looking in a $r \times r$ grid for the Z-plane that has an amount of bright pixels larger than a certain threshold. Despite the presence of a bright uniform cuticle layer above the cell layer, *PreMosa* still selects the latter for projection with an adequate set of parameters. This is probably due to the fact that the pixels in the membranes of the cell are themselves slightly brighter than the cuticle pixels, and because *PreMosa* offers a step where the surface is regularized. Indeed, *PreMosa* performs very well in areas of the sample where the layer is quasi-flat. However defects appear in regions where the curvature is higher (arrow), a fact also noted in [13] on other samples. As explained in [7], the grid size parameter must be significantly smaller than the cross-section of the tissue layer with any XY plane. In its most curved section, this cross section can be as small as 2 cell diameters, which is about 40-60 pixels in this region. In our tests the optimal grid-size value we found is 34, a value too large for this small cross section.

The *Extended Depth of Field* method was made to deal with transmitted light images. It is therefore not very well suited to our confocal test image. However we could find some parameter combination that extracts most of the tissue layer. Nonetheless, the height-map generated is very irregular. We can see the cells membrane in the projection, but their interior is filled with high and noisy signal. The error-map for this method has therefore high values everywhere.

Since we could configure the *Min Cost Z Surface* plugin to detect two surfaces, it reliably detected both the cuticle layer and the cell layer. The projection obtained with this second surface is satisfying. Like for the *LocalZProjector* projection, it however displays some small inaccuracies in its flat regions.

Along with the latter, *Fast SME* is the second energy-minimization based method of this comparison. It is built to specifically detect the layer of cells stained for their membrane, without the specification of any parameter. It succeeds to do so, except in one region (arrow) where the height-map has a jump to the cuticle layer in the region where it is bright. Here the cells are the largest in the tissue. It is likely that at these locations, the pixel columns that go through the inside of the cells get incorrectly classified as foreground, because of the bright intensity of the cuticle layer. The reference-surface then deforms to reach out the cuticle layer at these positions.

**C. Comparing optimal height-maps.** Several methods tested can also return the height-map that encodes the reference surface used for the projection. The height-map image stores the index of the Z-plane for each (X, Y) position in the reference surface. The comparison of the resulting height-maps confirms the observations and differences between methods noted above (Figure S11).

The height-map outputs of *LocalZProjector*, *Stack Focuser* and *PreMosa* resemble each other closely. In the case of *PreMosa*, the height-map has patches where the difference with the ground-truth is 1 or larger in the left part of the image, where the curvature is large. The *Stack Focuser* method also displays a similar defect in regions of even higher layer curvature (bottom left part). More differences in RMSE arise from the small discontinuity patches caused by the signal of dead cells deep in the tissue, for high values of Z.

The height-map of the *Extended Depth of Field* method reveals the main reason for the large value of RMSE in the projection. The height-map lacks some larger-scale regularization that would bridge correct Z-plane values over the center of cells in the cell layer.

Contrary to the first three methods, the height-map returned by the *Min Cost Z Surface* plugin does not appear as large patches of constant integer values. The Z-values in this height-map are decimal, a feature we attribute to the original formulation of the problem in this method. These values vary locally at a small scale, which generates the texture we observe in the error-map for this height-map. Nonetheless, non-integer values are rounded when projecting the source stack, and despite the error in the height-map, the projection returned by the *Min Cost Z Surface* is satisfying (Figure S10).

Finally, we retrieve in the height-map returned by the *FastSME* method the large inexact patch where the reference surface jumps towards the cuticle layer.

### Supplementary Note 3: Comparing segmentation results on optimal projections.

The metrics reported in the Supplemental note 2B show how close can a projection result be from a ground truth built by manually annotating a tissue surface. Most of the time, the projection is meant to be used subsequently in the next part of an analysis pipeline, *e.g.* for cell segmentation. To further assess the performance of the *LocalZProjector*, we investigated what is the impact of the projection method on cell segmentation accuracy. To do so, we run a simple cell segmentation pipeline on all the optimal projections obtained above, including the ground-truth projection, and compare results to a segmentation ground truth obtained manually. The results of this comparison are plotted in Figure S3d.

**A. The object-level consistency error metric.** In our case the segmentation results of the *Drosophila* dataset used in this work consists in about 3000 simply-connected regions corresponding each to a binary mask of a cell in the tissue layer. There exists metrics to measure the accuracy of such a segmentation result against a ground truth. In [14] the authors propose the use of the amount of overlap between the ground-truth and computed masks, averaged over all cells. While simple and intuitive, this metric tends to minimize the negative impact of over and under-segmentation, which are the two major defects that strongly affect segmentation of tissue images where cells are stained for their membrane. Instead we propose the use of the object-consistency error, introduced in [15], specifically built to highlight the impact of under and over-segmentation.

The authors first define a partial error measure $\mathrm{E}_{g,s}$ for two segmentation results $I_g$ and $I_s$. $I_g$ is made of $M$ non-overlapping masks $A_j$, and $I_s$ is made of $N$ non-overlapping masks $B_i$. This partial error measure is the average of an error measure $\mathrm{e}_j$ for each mask $A_j$ weighted by a factor $W_j$:

$$\mathrm{E}_{g,s}(I_g, I_s) = \sum_{j=1}^{M} \mathrm{e}_j \times W_j \tag{2}$$

The weights $W_j$ are the area proportion of the mask $A_j$:

$$W_j = \frac{|A_j|}{\sum\limits_{l=1}^{M} |A_l|} \tag{3}$$

where $|.|$ denotes the cardinal of a mask. The error $\mathrm{e}_j$ for the mask $A_j$ is equal to 1 minus the intersection over union weighted by a factor $W_{ji}$.

$$\mathrm{e}_j = 1 - \sum_{i=1}^{N} \frac{|A_j \cap B_i|}{|A_j \cup B_i|} \times W_{ji} \tag{4}$$

where $W_{ji}$ is given by:

$$W_{ji} = \frac{\bar{\delta}_{ji} \times |B_i|}{\sum_{k=1}^{N} \bar{\delta}_{jk} \times |B_k|} \quad \textbf{(5)}$$

$$\bar{\delta}_{ji} = \begin{cases} 1, & \text{if the intersection of} A_j \text{and} B_i \text{ is not empty} \\ 0, & \text{otherwise} \end{cases} \quad \textbf{(6)}$$

Finally, the object-consistency error $oce$ is calculated as $oce = \min(\mathrm{E}_{g,s}, \mathrm{E}_{s,g})$. Even within a set made of a rather large number of masks, the object-consistency error reliably penalizes over-segmentation or false-splits and under-segmentation or false-merges, respectively where a cell is split over several disjoint masks and where a segmentation mask covers several cells. We implemented this metric in MATLAB and used its results in Figure S3d and Table 1 in the main text.

**B. Simple segmentation pipeline.** From the optimal projections obtained previously, we run a straightforward, fully automated segmentation pipeline in Fiji [3], that returns a binary mask where each cell is represented by a connected set of white pixels separated by a black ridge. The pipeline is as follow:

- Filter the projection with a 3x3 median filter.
- Filter the resulting image with a Gaussian filter with $\sigma = 0.7$ pixels.
- From the MorphoLibJ [16] package in Fiji, run the Morphological Segmentation tool with a tolerance of 120 and a connectivity of 4.

We then obtain directly the label image $I_s$ we can use to measure the $oce$ value. This simplistic pipeline is not expected to fare perfectly against the amount of noise and staining imperfections in the test image but it is however good enough to demonstrate the impact of a projection method on the downstream analysis results.

**C. Ground-truth for segmentation accuracy.** We generated the segmentation ground-truth image $I_g$ by running the pipeline above on the ground-truth projection described in the Supplemental note 1. The numerous remaining segmentation errors were then manually corrected in Fiji.

| | RMSE projection | RMSE height map | Timing (s) | OCE segmentation |
|---|---|---|---|---|
| *MIP* | 180 | | 0.05 | 0.350 |
| *StackFocuser* | 96.7 | 71.1 | 5.17 | 0.141 |
| *SurfCut* | 171 | | 1.50 | 0.581 |
| *PreMosa* | 79.9 | 1.38 | 1.60 | 0.100 |
| *EDF* | 146 | 3.59 | 6.91 | 0.261 |
| *MinCostZ* | 138 | 2.15 | 8.7 | 0.290 |
| *FastSME* | 81.5 | 0.49 | 10.0 | 0.078 |
| *LocalZProjector* | 76.8 | 1.18 | 4.96 | 0.080 |
| *LocalZProjector 2nd best* | 76.9 | 1.08 | 0.35 | 0.066 |

**Table S1.** Performance comparison on a high quality image of a *Drosophila* pupa notum, captured with a laser-scanning confocal microscope (LSM 880, Zeiss). This specific image is of high quality compared to that used for the comparison reported in Table 1. There are little spurious structures, the cuticle auto-fluorescence signal is faint and there are almost no dead bodies. All tools give visually satisfactory projection results, but the quality of the projection strongly impacts the segmentation accuracy. For *LocalZProjector*, we include metrics for a second parameter set with a larger binning value, resulting in a ten times quicker projection with only a minor cost on the projection RMSE. The raw data and the methodology and tools to perform this comparison are available online [17].

| | RMSE projection | RMSE height map | Timing (s) | OCE segmentation |
|---|---|---|---|---|
| *MIP* | 4750 | | 0.5 | 0.229 |
| *StackFocuser* | 2130 | 8.26 | 11.2 | 0.327 |
| *SurfCut* | 1080 | | 24 | 0.229 |
| *PreMosa* | 1730 | 4.89 | 180 | 0.288 |
| *EDF* | 2030 | 8.02 | 84.2 | 0.311 |
| *MinCostZ* | 2630 | 2.08 | 178 | 0.211 |
| *FastSME* | 1370 | 4.54 | 87.8 | 0.215 |
| *LocalZProjector* | 944 | 2.22 | 5.3 | 0.162 |

**Table S2.** Performance comparison on the adult zebrafish brain image used in Figure 5 of the main text. This image is a rather large compared to the image used in Table S1 (551 Gpixels *vs* 29 Gpixels) and we see the impact on the processing time. The raw image is available online [18].

### List of Figures

S1 Overview of the Local Z Projector method. The source 3D stack is first processed slice by slice. Each 2D slice is binned to accelerate processing, denoised with a Gaussian filter, then the main filter is applied. The main filter is chosen in type (mean filter, standard deviation filter, ...) and in size to have a strong response in the layer of interest. The z plane for which this response is maximal is stored for every (X, Y) position in the height-map image. The height-map is then median-filtered, to remove local spurious irregularities, and up-scaled back so that its width and height match those of the source 3D stack. Finally, the height-map is used as a reference to read pixels in the source stack around the $z$ position it contains, accumulating $\pm\Delta z$ planes around this reference plane. For a given main filter type and accumulation method, four parameters must be specified: the binning, the sigma of the Gaussian filter, the size of the main filter and the value of $\Delta z$. . . . . . . . . . . . . . . . . . . 13

S2 Manual generation of a ground truth dataset. From left to right: small region in four consecutive slices in the 3D dataset, annotated in green for regions where cells of interest are in focus. . . . . . . . . . . . . . . . . . . 14

S3 Performance metrics comparison for projection tools. In the four plots, lower is better. **a.** Root mean square error (RMSE) comparing the projection obtained with several methods against a ground-truth projection. **b.** RMSE for the comparison of the height-map yielded by the same methods. As the MIP and SurfCut methods do not return a height-map image, they are not included in this plot. **c.** Time required to perform the projection, taken as the median of 5 runs. **d.** Object-consistency error (OCE) for cell segmentation results on the projection image generated by each image. . . . . . . . . . . . . . . . . . . . . . . . . . . . . . . . . . . . . . . . . 15

S4 Optimization of the parameters of the *Local Z Projector* method. Except for the parameter varied in each plot, the plugin was run with the following parameters, corresponding to a minimum in RMSE value: binning = 2, Gaussian sigma = 0.5, standard-deviation filter size = 13, median filter half-size = 16, $\Delta z = 1$, and accumulating by taking the brightest pixel. The blue curve gives the RMSE value (Equation 1) and the orange curve gives the execution time in seconds of the projection on the test computer (see Supplemental methods). . . . . . . . . . . 16

S5 Optimization of the filter-size parameter of the *Stack Focuser* method. . . . . . . . . . . . . . . . . . . . . . . 17

S6 Optimization for the parameters of the *PreMosa* method. Except for the parameter varied in each plot, the tool was run with the following parameters, corresponding to a minimum in RMSE value: Grid-size = 34, Distance = 1, Bright-pixel threshold = 20%. . . . . . . . . . . . . . . . . . . . . . . . . . . . . . . . . . . . . . 18

S7 Optimization for the parameters of the *Extended Depth of Field* method. Each point correspond to a combination of a quality parameter set and a topology parameter set, taken from the simple version of the tool user-interface. The median running time for a given quality parameter set over all topology parameter set is given in orange as annotation on the plot. Topology parameters: A. No smoothing, B. Median 3 × 3, C. Median 3 × 3 + Gaussian σ=1, D. Median 3 × 3 + Gaussian σ=2 + Closing, E. Median 5 × 5 + Gaussian σ=4 + Opening + Closing. Quality parameters: MCC stands for Majority Consistency Checks. . . . . . . . . . . . . . . . . . 19

S8 Optimization for the parameters of the *Minimum Cost Z Surface* method. Each point correspond to a parameter set determined empirically. Unless specified, the sampling parameters required by the tool are always $d_{xy} = 0.25$ and $dz = 1$, corresponding to downsampling in X and Y by a factor of 4 (except for the leftmost point for which is $d_{xy} = 0.5$), Maximal distance between surfaces = 15, Minimal distance between surfaces = 3, Maximal delta z between adjacent voxels = 1. . . . . . . . . . . . . . . . . . . . . . . . . . . . . . . . . . . 20

S9 Optimization for the parameters of the *FastSME* method. . . . . . . . . . . . . . . . . . . . . . . . . . . 21

S10 Projection results comparison for 8 methods. The top line shows the ground-truth projection (1148 × 1148), the scale for projection image and the scale for error maps. The two bottom columns show the results of the projection by one method, next to the error map. The error map is obtained by measuring the pixel-by-pixel absolute difference with the ground-truth projection, filtered by a median of size 41 × 41. Yellow arrows refer to salient points described in the text. *MIP*: Maximum intensity projection. *E.D.F*: Extended depth of field. . . 22

S11 Height-map results comparison for 6 methods. The top line shows the ground-truth height-map, the color-scale matching the Z-plane in the source stack and the scale for error maps. The value 0 (black) in the ground-truth height-map corresponds to a location in the image where there is no cell layer. The two bottom columns show the results of the projection by one method, next to the error map. *E.D.F*: Extended depth of field. When possible, we use decimal values in the height-maps to generate this figure. . . . . . . . . . . . . . . . . . . 23

S12 Qualitative performance of *LocalZProjector* on other datasets. The 3D images used for this figure are taken from [13]. Using the LocalZProjector tool, we derived parameters that would give a projection similar to what is presented in Figure 6 of [13]. . . . . . . . . . . . . . . . . . . . . . . . . . . . . . . . . . . . . . . . . . 24

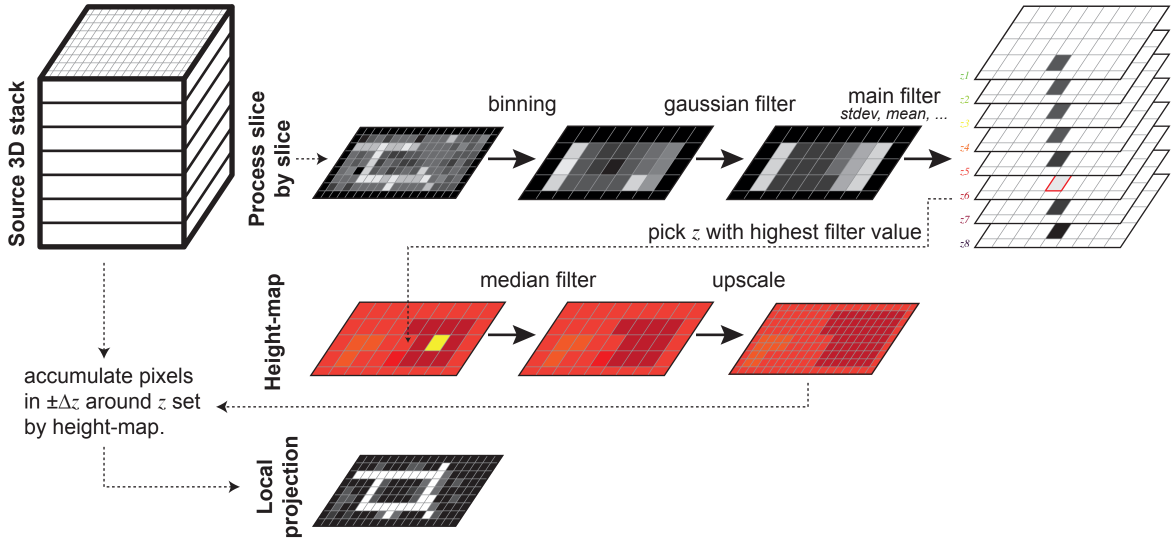

**Figure S1.** Overview of the Local Z Projector method. The source 3D stack is first processed slice by slice. Each 2D slice is binned to accelerate processing, denoised with a Gaussian filter, then the main filter is applied. The main filter is chosen in type (mean filter, standard deviation filter, ...) and in size to have a strong response in the layer of interest. The z plane for which this response is maximal is stored for every (X, Y) position in the height-map image. The height-map is then median-filtered, to remove local spurious irregularities, and up-scaled back so that its width and height match those of the source 3D stack. Finally, the height-map is used as a reference to read pixels in the source stack around the $z$ position it contains, accumulating $\pm\Delta z$ planes around this reference plane. For a given main filter type and accumulation method, four parameters must be specified: the binning, the sigma of the Gaussian filter, the size of the main filter and the value of $\Delta z$.

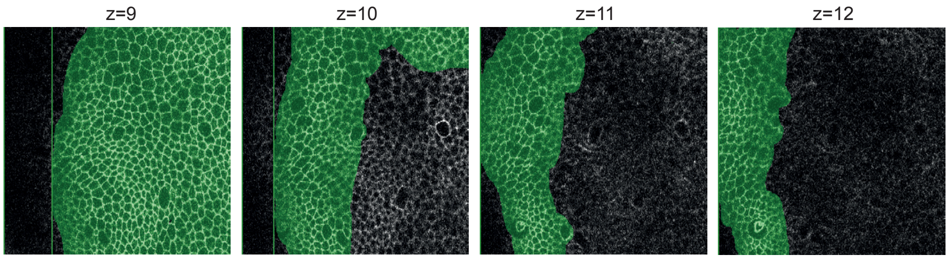

**Figure S2.** Manual generation of a ground truth dataset. From left to right: small region in four consecutive slices in the 3D dataset, annotated in green for regions where cells of interest are in focus.

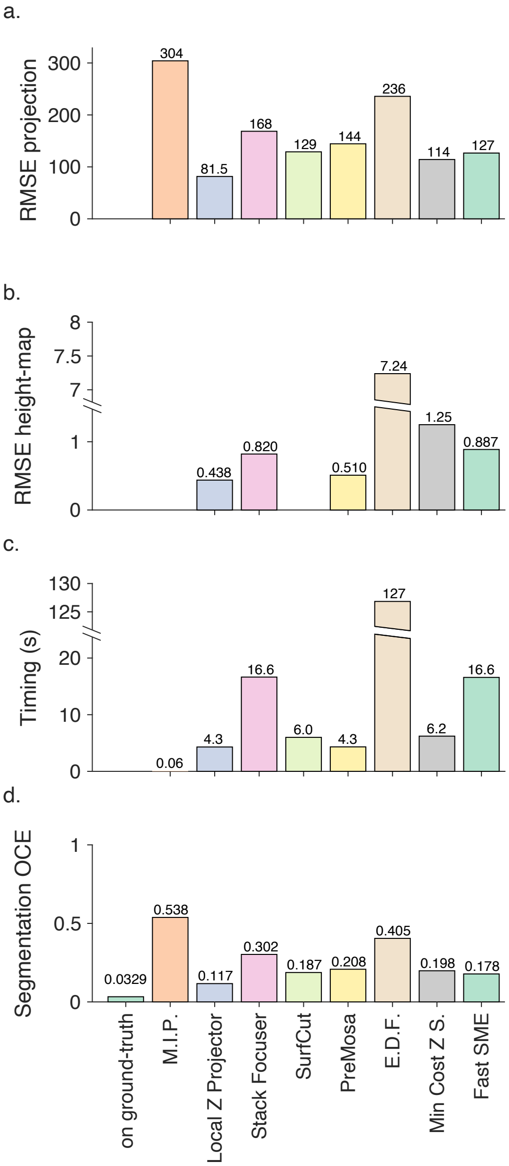

**Figure S3.** Performance metrics comparison for projection tools. In the four plots, lower is better. **a.** Root mean square error (RMSE) comparing the projection obtained with several methods against a ground-truth projection. **b.** RMSE for the comparison of the height-map yielded by the same methods. As the MIP and SurfCut methods do not return a height-map image, they are not included in this plot. **c.** Time required to perform the projection, taken as the median of 5 runs. **d.** Object-consistency error (OCE) for cell segmentation results on the projection image generated by each image.

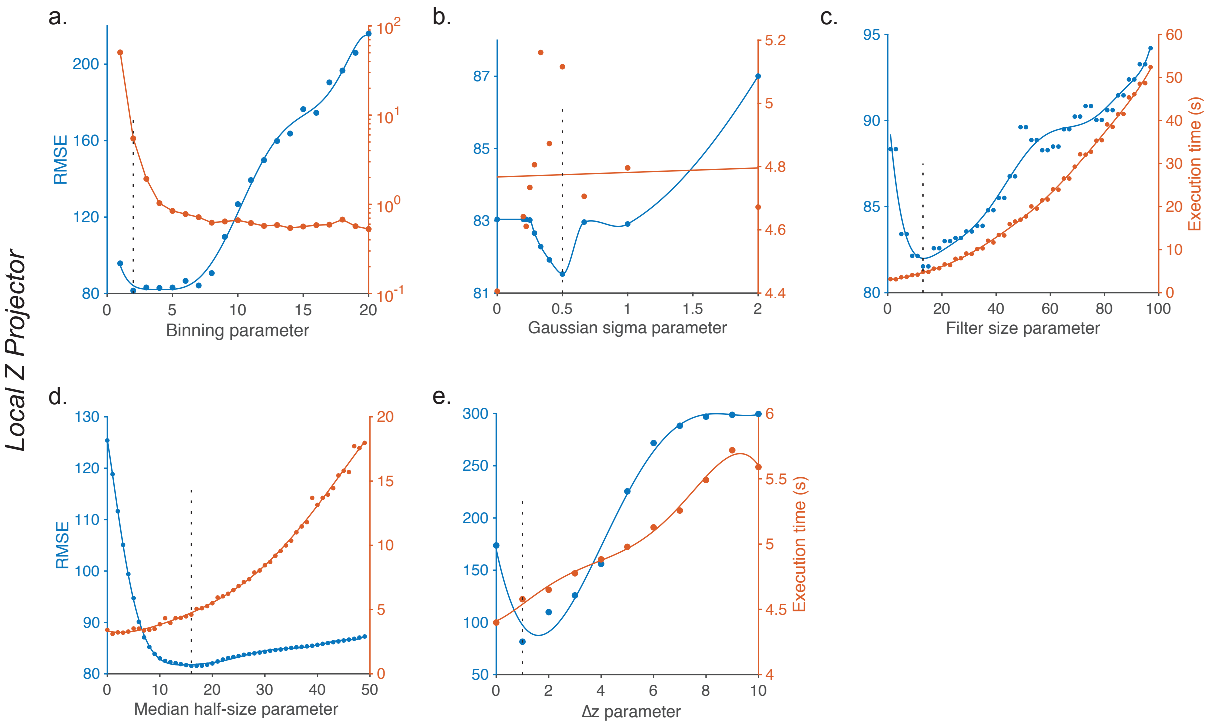

**Figure S4.** Optimization of the parameters of the *Local Z Projector* method. Except for the parameter varied in each plot, the plugin was run with the following parameters, corresponding to a minimum in RMSE value: binning = 2, Gaussian sigma = 0.5, standard-deviation filter size = 13, median filter half-size = 16, $\Delta z = 1$, and accumulating by taking the brightest pixel. The blue curve gives the RMSE value (Equation 1) and the orange curve gives the execution time in seconds of the projection on the test computer (see Supplemental methods).

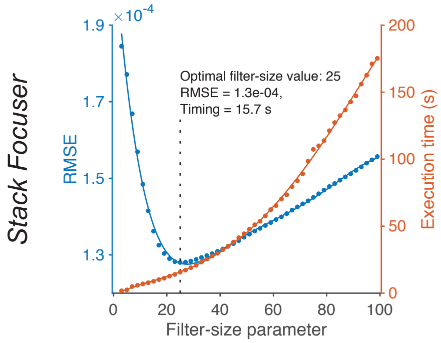

**Figure S5.** Optimization of the filter-size parameter of the *Stack Focuser* method.

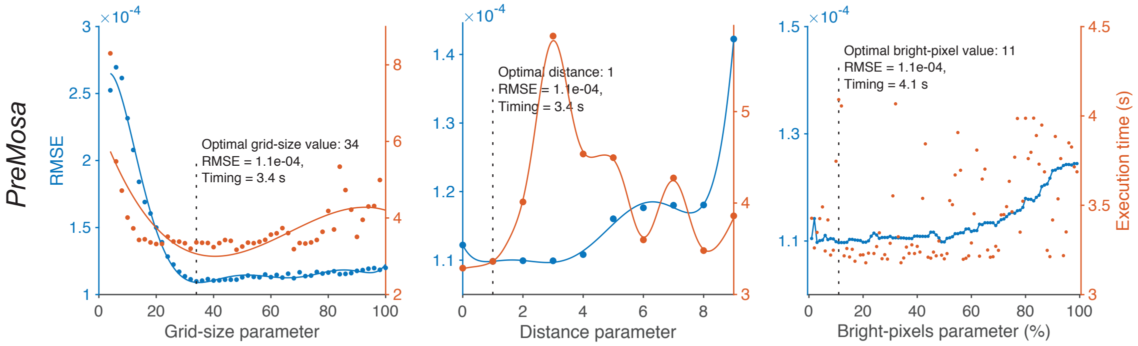

**Figure S6.** Optimization for the parameters of the *PreMosa* method. Except for the parameter varied in each plot, the tool was run with the following parameters, corresponding to a minimum in RMSE value: Grid-size = 34, Distance = 1, Bright-pixel threshold = 20%.

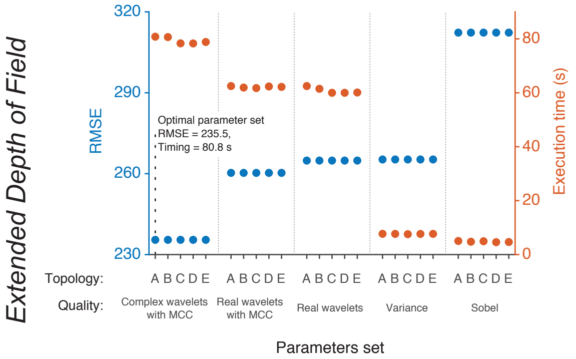

**Figure S7.** Optimization for the parameters of the *Extended Depth of Field* method. Each point correspond to a combination of a quality parameter set and a topology parameter set, taken from the simple version of the tool user-interface. The median running time for a given quality parameter set over all topology parameter set is given in orange as annotation on the plot. Topology parameters: A. No smoothing, B. Median 3 × 3, C. Median 3 × 3 + Gaussian σ=1, D. Median 3 × 3 + Gaussian σ=2 + Closing, E. Median 5 × 5 + Gaussian σ=4 + Opening + Closing. Quality parameters: MCC stands for Majority Consistency Checks.

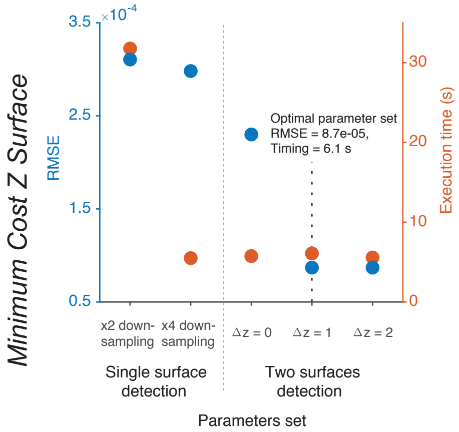

**Figure S8.** Optimization for the parameters of the *Minimum Cost Z Surface* method. Each point correspond to a parameter set determined empirically. Unless specified, the sampling parameters required by the tool are always $d_{xy} = 0.25$ and $dz = 1$, corresponding to downsampling in X and Y by a factor of 4 (except for the leftmost point for which is $d_{xy} = 0.5$), Maximal distance between surfaces = 15, Minimal distance between surfaces = 3, Maximal delta z between adjacent voxels = 1.

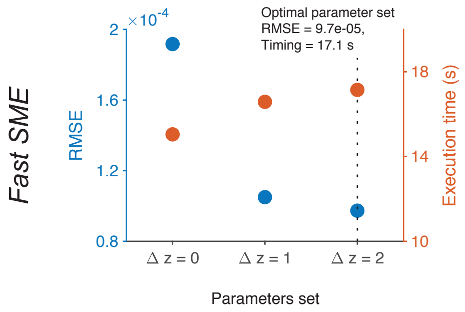

**Figure S9.** Optimization for the parameters of the *FastSME* method.

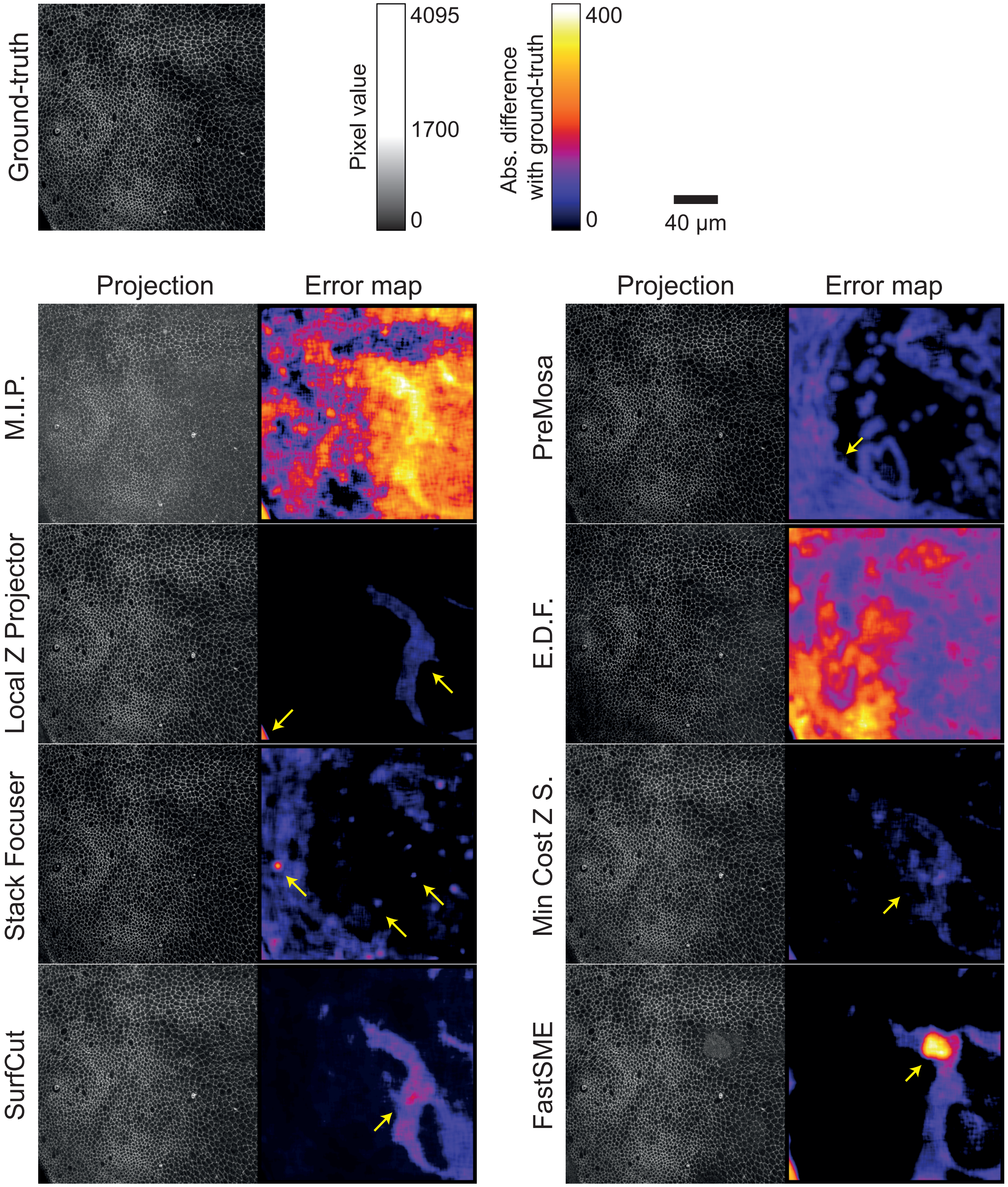

**Figure S10.** Projection results comparison for 8 methods. The top line shows the ground-truth projection (1148 × 1148), the scale for projection image and the scale for error maps. The two bottom columns show the results of the projection by one method, next to the error map. The error map is obtained by measuring the pixel-by-pixel absolute difference with the ground-truth projection, filtered by a median of size 41 × 41. Yellow arrows refer to salient points described in the text. *MIP*: Maximum intensity projection. *E.D.F*: Extended depth of field.

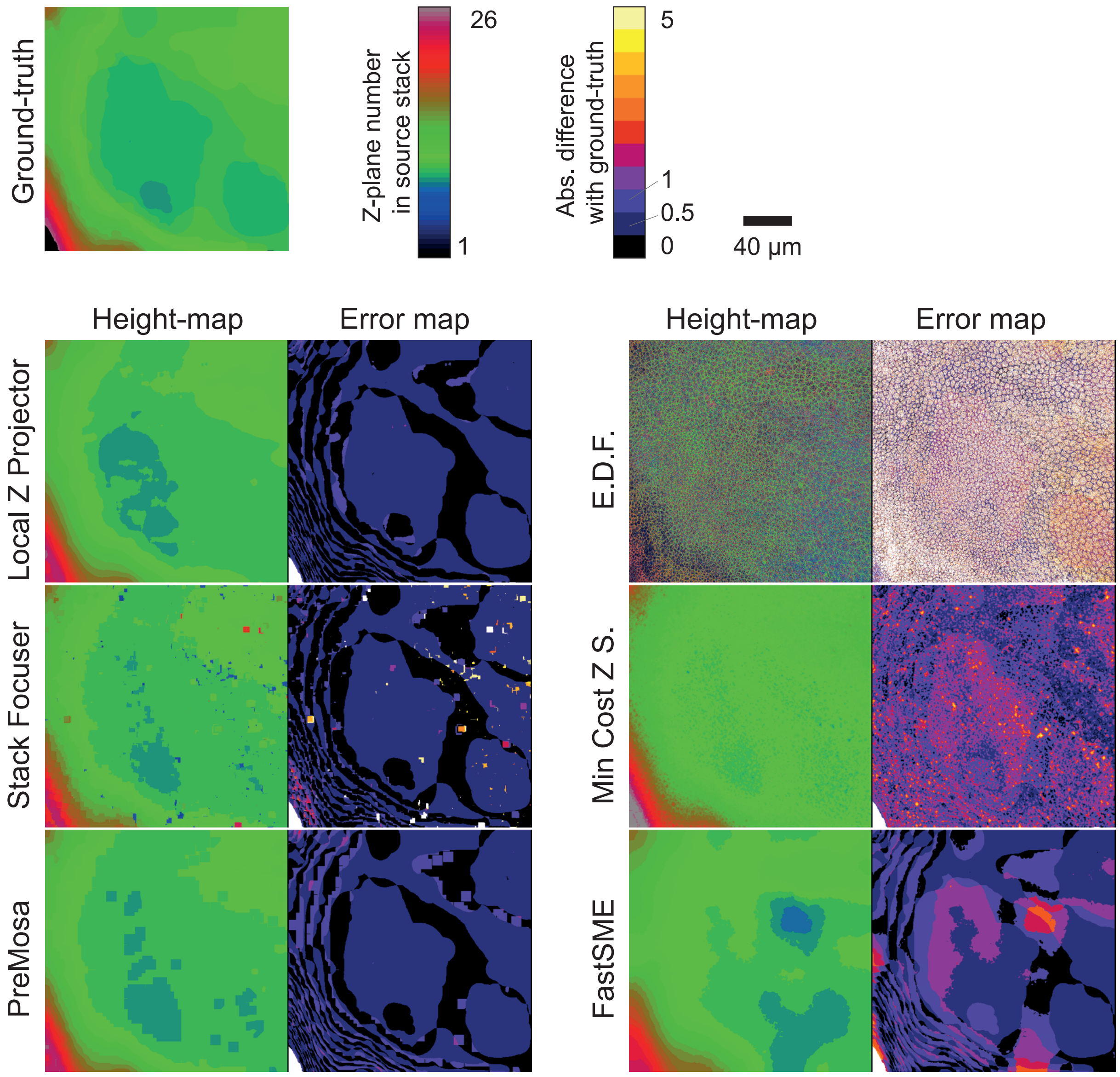

**Figure S11.** Height-map results comparison for 6 methods. The top line shows the ground-truth height-map, the color-scale matching the Z-plane in the source stack and the scale for error maps. The value 0 (black) in the ground-truth height-map corresponds to a location in the image where there is no cell layer. The two bottom columns show the results of the projection by one method, next to the error map. *E.D.F*: Extended depth of field. When possible, we use decimal values in the height-maps to generate this figure.

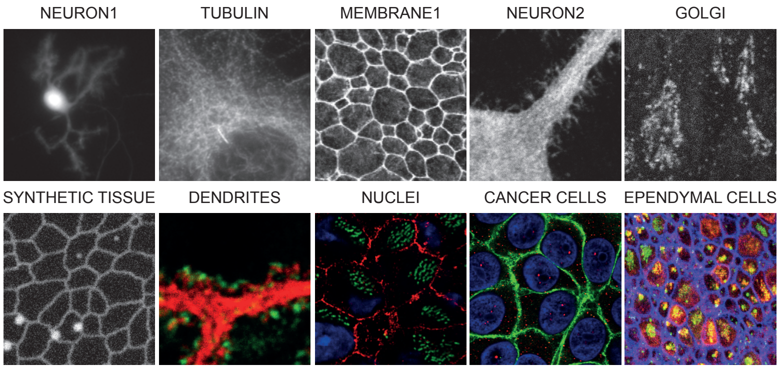

**Figure S12.** Qualitative performance of *LocalZProjector* on other datasets. The 3D images used for this figure are taken from [13]. Using the LocalZProjector tool, we derived parameters that would give a projection similar to what is presented in Figure 6 of [13].
